# ACTL6A regulates the Warburg effect through coordinated activation of AP-1 signaling in head and neck squamous cell carcinoma

**DOI:** 10.1101/2025.09.02.671886

**Authors:** Mehri Monavarian, Keene Lee, Alistaire R. Sherman, Imran A. Mohammad, Sainiteesh Maddineni, Mao Zhang, Joseph C. Wu, Katrin F. Chua, John Sunwoo, Andrey Finegersh

## Abstract

Tumors adapt to hypoxic environments by activating stress-responsive transcriptional programs; however, the chromatin structure permitting these adaptations is poorly defined. We identify ACTL6A, a SWI/SNF subunit over-expressed in several cancers, as a critical regulator of enhancer architecture that drives glycolysis and hypoxia gene expression in head and neck squamous cell carcinoma. ACTL6A expression coordinates AP-1 chromatin accessibility at enhancers while simultaneously amplifying MAPK/ERK signaling through Ras and Galectin-1. ACTL6A is required for the metabolic effects of MEK inhibitors and is strongly correlated with induction of p-ERK and MAPK signaling programs across human cancers. Depleting ACTL6A collapses hypoxic adaptation *in vivo* and inhibits glycolysis, leading to increased mitochondrial dependency. Using spatial biology, we define an ACTL6A-dependent transcriptional signature in human tumors that drives hypoxia and MAPK signaling and marks a spatially-isolated malignant cell niche. ACTL6A depletion induces a metabolic vulnerability in treatment resistant tumors that sensitizes them to IACS-010759 *in vivo*. These results implicate ACTL6A as an epigenetic switch that sustains a chromatin architecture permissive for hypoxic adaptation and metabolic plasticity, which can be exploited to target SWI/SNF-amplified tumors.

## INTRODUCTION

Cancer cells shift their metabolism toward aerobic glycolysis, which promotes hypoxic tumor growth, immune evasion, and treatment resistance (1,2). Shunting metabolism from oxidative phosphorylation also contributes to cell survival and growth by decreasing production of reactive oxygen species (ROS) and generating glycolytic metabolites for anabolic pathways (3). While this shift was originally described by Otto Warburg in the 1920s (4), the regulatory mechanisms that rewire cancer cell metabolism remain poorly understood. Importantly, defining epigenetic drivers that generate a chromatin structure permissive for metabolic plasticity may identify vulnerabilities that could be therapeutically targeted.

Copy number amplification of chromosome 3q26 occurs in many cancers and up to 75% of squamous cell carcinomas (SCC) (5), where it is associated with over-expression of several oncogenic drivers (6). One of these oncogenes, ACTL6A, is a component of the SWI/SNF (BAF) complex, which is an ATP-dependent chromatin remodeling complex that is critical in the development and maintenance of cell fate (7), RNA polymerase II mediated gene transcription (8), and DNA repair (9). ACTL6A stoichiometrically regulates SWI/SNF complex occupancy in SCC (10), where it drives TEAD/YAP activation to maintain a poorly differentiated cancer cell phenotype (10–12). Recent studies in other cancers have also linked ACTL6A expression with maintenance of metabolic phenotypes, including activation of glutathione synthesis and inhibition of ferroptosis in gastric carcinoma (13) and stimulation of glycolysis in ovarian cancer (14); however, ACTL6A chromatin occupancy has not been directly defined and specific mechanisms by which it generates metabolic transcriptional programs are unknown.

Head and neck squamous cell carcinoma (HNSCC) is the sixth most common malignancy (15) and is one of the few cancers with declining overall survival and increasing incidence in North America (16), suggesting an urgent need to develop treatments for this malignancy. Like other solid tumors, HNSCC shows increased dependence on glycolysis (17), with increasing rates of glycolysis in tumors leading to worse survival and decreased immune cell infiltration (18). Therefore, identifying and targeting mechanisms underlying metabolic plasticity in this cancer has the potential to improve treatment outcomes.

Using combined epigenomic, single-cell spatial transcriptomic, and metabolic functional assay approaches, we identify ACTL6A expression as a molecular switch that drives aerobic glycolysis and hypoxic adaptation in HNSCC. We define concurrent mechanisms by which ACTL6A regulates AP-1-mediated transcriptional programs through regulation of chromatin accessibility at glycolysis and hypoxia gene enhancers as well as potentiating upstream signaling through Ras/LGALS1. By establishing ACTL6A as a key regulator of metabolic plasticity in HNSCC, we develop a synthetic lethality strategy to drive tumor killing in a treatment resistant oral cavity cell line. These results define a key epigenetic mechanism that drives the Warburg effect in solid tumors and a novel strategy to overcome treatment resistance.

## RESULTS

### ACTL6A levels regulate aerobic glycolysis in HPV (-) HNSCC

Within The Cancer Genome Atlas (TCGA) HNSCC cohort, ACTL6A over-expression significantly reduces overall survival in HPV (-) but not HPV (+) HNSCC (Fig. 1A) (19). As HPV (-) HNSCCs have higher lethality (20) and a survival association with ACTL6A, we chose to focus on this cancer subset. We then performed a correlational analysis on 62 cell lines derived from HPV (-) oral cavity and larynx SCC from DepMap to assess if glycolysis gene expression is correlated with ACTL6A levels across different mutational backgrounds (21). We found significant positive correlations between ACTL6A and expression of all glycolysis genes (Fig. 1B and table S3). Based on this, we performed RNA-seq on the HPV (-) HNSCC cell line SCC1 following transient ACTL6A knockdown (KD) (fig. S1A) with control and KD replicates separating cleanly by PCA (fig. S1B). ACTL6A depletion significantly reduced expression of the glycolytic enzymes ENO2, GAPDH, PKM, PFKP, and PGK1 (Fig. 1C). Gene set enrichment (GSEA) using the MSigDB Hallmark gene set showed significant down-regulation of hypoxia, glycolysis, and KRAS pathways in the ACTL6A depleted samples (Fig. 1D).

**Fig. 1.**
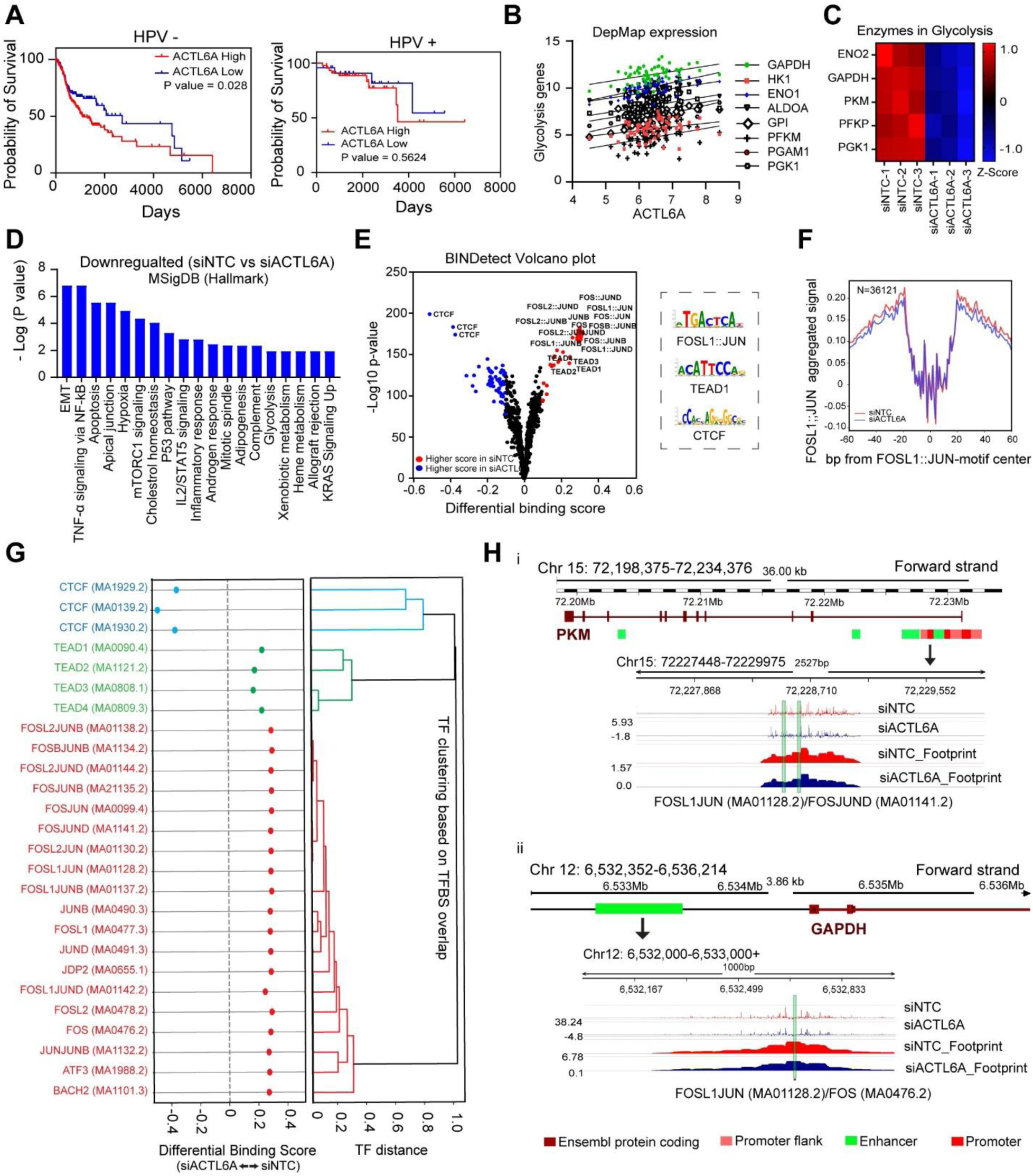
ACTL6A is associated with increased aerobic glycolysis in HNSCC and regulates AP-1 transcription factor chromatin accessibility. (A) Survival analysis using log-rank test in HPV (-) (left) and HPV (+) (right) HNSCCs from TCGA (B) Correlations between ACTL6A and glycolysis gene expression in 62 HPV (-) HNSCC cell lines annotated in DepMap. (C) Heatmap of glycolysis gene expression in siACTL6A and control SCC1 cells using Z-scores calculated from RNA-seq data. (D) Most significantly down-regulated Hallmark pathways in SCC1 cells after siACTL6A. (E) Volcano plot from BINDetect analysis using TOBIAS tool on ATAC-seq data from SCC1 cells showing differential binding scores for transcription factors (TF) motifs in control samples compared with ACTL6A knockdown. (F) Aggregated footprinting plot, comparing SCC1 siNTC and siACTL6A samples, centered on predicted binding sites for FOSL1/JUN. (G) Differential binding scores for indicated transcription factors in top three clusters (shown separately). TF clustering is based on overlapping of TF binding sites. (H) Genomic tracks for PKM (i) and GAPDH (ii) were made using GRCh38.p14 assembly in Ensembl with their corresponding corrected ATAC-seq peaks and footprints generated with TOBIAS from SCC1 ATAC-seq data comparing the control samples with siACTL6A.

### ACTL6A modulates chromatin accessibility at AP-1 transcription factor sites

To assess the role of ACTL6A expression on transcription factor (TF) binding and identify mechanisms contributing to our metabolic phenotype, we performed ATAC-seq (assay for transposase-accessible chromatin) using SCC1 cells following transient ACTL6A knockdown (fig. S1). Differential accessibility analysis showed reduced accessibility in 3176 regions and increased accessibility in 1250 regions in ACTL6A knockdown cells compared with control. Then, we performed footprinting analysis using TOBIAS (22) to predict differential TF binding and clustering. Using motifs from JASPAR (23), we found three main clusters with differential activity following ACTL6A depletion, with FOSL1::JUN (AP-1) and TEAD transcription factor clusters showing significantly increased chromatin accessibility in the siNTC samples and CCCTC-binding factor (CTCF) cluster with increased chromatin accessibility in siACTL6A samples (Fig. 1E,G and fig. S1D). To further visualize the difference in binding activity across all binding sites between ACTL6A depletion and control cells, aggregated signal was plotted around motif center, which confirmed reduced accessibility around AP-1 and TEAD1 motifs (Fig. 1F fig. S1E). We conducted pathway analysis using MSigDB Hallmark on 568 overlapping genes with less accessible regions on ATAC-seq and downregulation on RNA-seq in siACTL6A samples. We found downregulation of hypoxia, KRAS signaling, and Glycolysis pathways (fig. S1). We then investigated the regulatory regions of glycolysis genes that were significantly down-regulated with siACTL6A and found reduced activity of the FOSL1::JUN TFs on the promoter and enhancer of PKM and GAPDH, respectively (Fig. 1H). These results indicate that ACTL6A expression maintains chromatin accessibility at AP-1 TF sites, which potentiate expression of genes involved in glycolysis.

### ACTL6A regulates chromatin accessibility of AP-1 sites in a SWI/SNF dependent manner

We performed CUT&RUN sequencing using an ACTL6A antibody on SCC1 cells following 72 hours treatment with either DMSO or 1µM of AU-15330, a PROTAC degrader of the SWI/SNF subunits SMARCA2 and SMARCA4 (fig. S2). AU-15330 resulted in significant depletion of ACTL6A occupancy (Fig. 2A). Loss of ACTL6A occupied peaks was concentrated at inter- and intra-genic regions consistent with an effect on enhancers as the residual ACTL6A peaks remaining after AU-15330 were redistributed toward promoters (31.6% to 53.1% of peaks) (Fig. 2B) (24). Motif enrichment analysis (HOMER) identified AP-1 family motifs (Fos, JunB, Fra1, Fra2, ATF3, and AP-1) as the most significantly enriched motifs in DMSO-treated cells, while YY1, a component of the INO80 chromatin remodeling complex, was most significantly enriched after AU-15330 treatment (Fig 2C). As ACTL6A is also a component of INO80 (25), this offers insights into its SWI/SNF independent genomic occupancy. To identify genes regulated by ACTL6A in a SWI/SNF dependent manner, we integrated RNA, ATAC, and CUT&RUN to reveal 265 genes intersecting across these datasets (Fig. 2D). GSEA analysis of these intersecting genes showed Glycolysis and Hypoxia among the top 10 most enriched pathways, along with other oncogenic pathways (Fig. 2E). Integrative analysis of ATAC and RNA-seq after ACTL6A depletion also showed that ACTL6A regulates expression of several oncogenes, including LGALS1 (Galectin-1) and ICAM1 (fig. S2).

**Figure 2.**
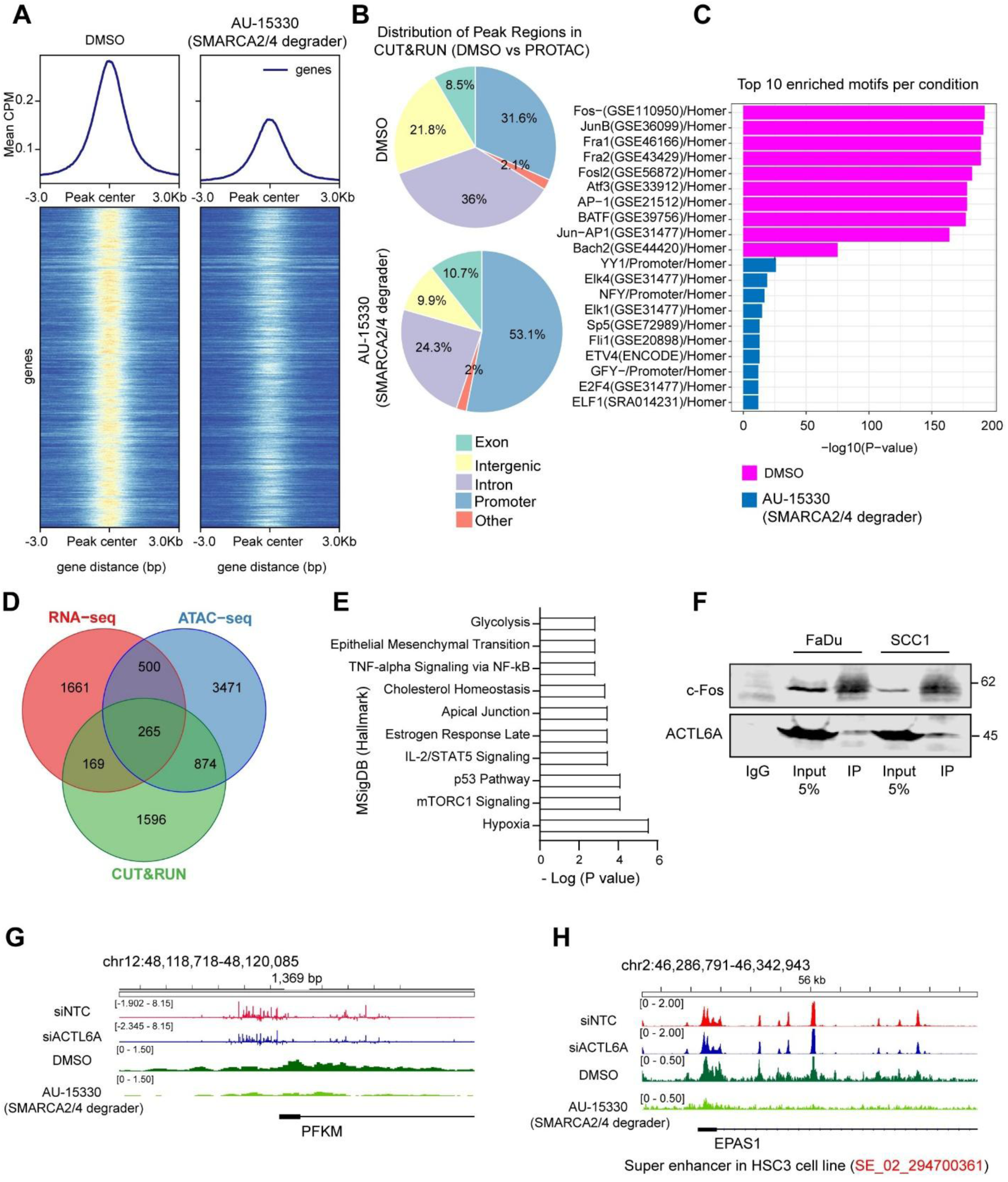
ACTL6A regulates chromatin accessibility of AP-1 transcription factor sites in a SWI/SNF-dependent manner. (A) Metagene plots (top) and heatmaps (bottom) centered on ACTL6A peaks (±3 kb) from ACTL6A CUT&RUN signal in SCC1 cells following 72 hours treatment with either DMSO or AU-1533. n=3/group (B) Pie charts show genomic distribution of ACTL6A CUT&RUN peaks in DMSO and AU-15330 treated SCC1 cells. SWI/SNF-dependent occupancy of ACTL6A is primarily at enhancers (intergenic and intronic) while ACTL6A maintains binding sites at promoters after SWI/SNF degradation. (C) Top enriched transcription factor motifs identified from ACTL6A CUT&RUN peaks in DMSO and AU-15330 treated cells using HOMER. (D) Venn diagram showing overlapping genes from RNA-, ATAC-, and CUT&RUN sequencing to identify those regulated by ACTL6A in SWI/SNF-dependent manner. (E) Top MSigDB Hallmark pathway analysis of 265 intersecting genes showing enrichment in hypoxia, glycolysis, and other oncogenic pathways. (F) Co-immunoprecipitation of ACTL6A and immunoblot for c-Fos and ACTL6A in FaDu and SCC1 cells; lanes show IgG control, 5% input, and c-FOS immunoprecipitate (IP). Integrative genomic browser tracks for the glycolytic enzyme (G) PFKM and (H) EPAS1/HIF2α showing ATAC-seq (siNTC, siACTL6A) and ACTL6A CUT&RUN (DMSO, AU-15330) signal. The EPAS1 locus overlaps a super-enhancer annotated in the HSC3 cell line (SE_02_294700361).

Because AP-1 motifs dominated ACTL6A-occupied regions, we tested whether ACTL6A physically associates with AP-1 transcription factors. Co-immunoprecipitation in native FaDu and SCC1 cells demonstrated that ACTL6A and c-Fos co-precipitate, with ACTL6A detected in the c-Fos immunoprecipitate but not in the IgG control (Fig. 2F). These data indicate that ACTL6A and c-Fos exist within common regions, providing a physical basis for ACTL6A regulation of AP-1 chromatin accessibility.

We then evaluated ACTL6A dependent regions within glycolysis and hypoxia genes, including PFKM, the rate limiting enzyme for glycolysis, and EPAS1 (HIF2α). These showed SWI/SNF-dependent ACTL6A enrichment that was abrogated with AU-15330 (Fig 2G,H). Interestingly, the EPAS1 region with differential binding for ACTL6A in CUT&RUN and ATAC-seq has been detected as a super enhancer in the HSC3 HPV-oral cavity SCC cell line (26) (SE_02_294700361, Fig 2H). Both of these genes had a much greater shift in intragenic chromatin accessibility and ACTL6A occupancy, consistent with an overall effect on enhancers.

### ACTL6A regulates hypoxic tumor growth and HIF-dependent transcriptional programs *in vitro* and *in vivo*

Integrative analyses of RNA-seq, ATAC-seq, and CUT&RUN sequencing data showed significant attenuation of AP-1 transcription factor accessibility at hypoxia and glycolysis related enhancers with ACTL6A depletion in HNSCC cells. We hypothesized that ACTL6A expression coordinates AP-1 dependent hypoxic adaptation in human cancers. We calculated sample-level AP-1 activity using DorothEA regulons and VIPER and correlated it with the Buffa hypoxia score across a TCGA cohort of solid tumors, stratifying each cohort into ACTL6A-high and ACTL6A-low tumors. In five cohorts (HNSCC, BRCA, OV, LIHC, STAD), AP-1 activity was significantly and positively correlated with hypoxia score in ACTL6A-high tumors (HNSCC r= 0.33, p < 0.001; BRCA r = 0.31, p < 0.001; OV r = 0.31, p = 0.0017; LIHC r = 0.36, p < 0.001; STAD r = 0.18, p = 0.033), whereas the correlation was weak and non-significant in ACTL6A-low tumors (r = 0.05, 0.09, 0.14, −0.13, and 0.12, respectively; all p > 0.05) (Fig. 3A). These data indicate that the coupling between AP-1 activity and the hypoxic response is dependent on high ACTL6A expression and is conserved across multiple malignancies.

**Figure 3.**
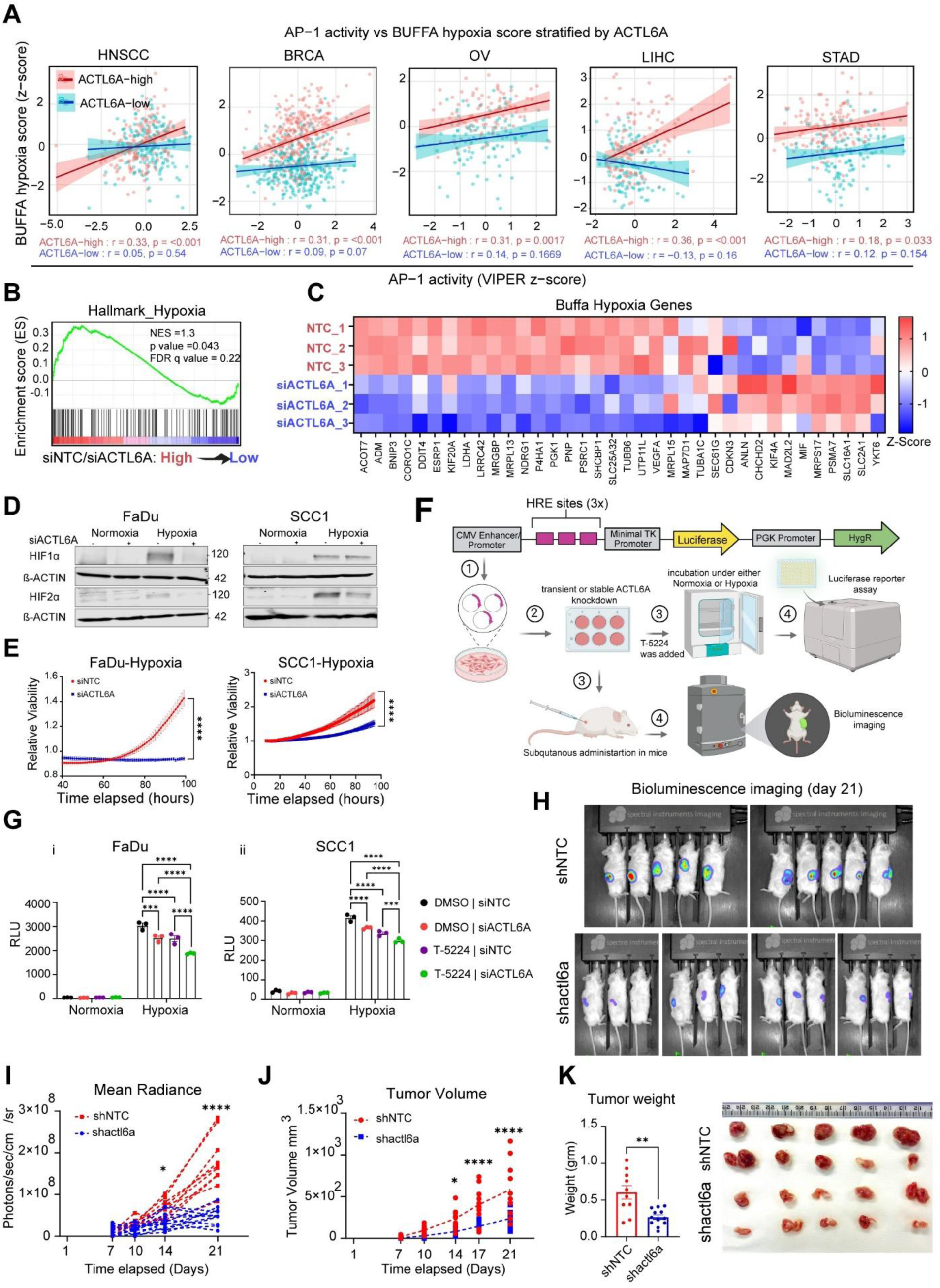
ACTL6A expression coordinates AP-1 activity with hypoxic tumor response and is required for HIF-dependent transcriptional programs *in vitro* and *in vivo*. (A) Correlation between AP-1 transcription factor activity (DorothEA regulons and VIPER z-score) and the Buffa hypoxia score across five TCGA cohorts (HNSCC, BRCA, OV, LIHC, STAD), with tumors stratified into ACTL6A-high (red) and ACTL6A-low (blue) groups. Spearman r and p values are shown for each group. (B) Enrichment plot for the Hallmark hypoxia gene set from SCC1 RNA-seq comparing control (siNTC) and ACTL6A-depleted (siACTL6A) cells. (C) Heatmap of Buffa hypoxia gene expression (Z-scores from RNA-seq) in siNTC and siACTL6A SCC1 cells (n = 3 per group). (D) Representative Western blots of HIF1α, HIF2α, and β-actin in FaDu (left) and SCC1 (right) cells following transient ACTL6A knockdown and 24-hour incubation in normoxia or 0.2% hypoxia. (E) Relative viability of FaDu (left) and SCC1 (right) cells following transient ACTL6A knockdown, measured under 0.2% hypoxia over the indicated time course using the Xcellbio AVATAR system. n = 6. **** p < 0.0001, two-way ANOVA followed by Sidak’s multiple comparisons test. (F) Schematic of the 3× HRE-luciferase reporter construct and the experimental workflow: reporter transduction, transient or stable ACTL6A knockdown, incubation in normoxia or hypoxia with DMSO or the AP-1 inhibitor T-5224, followed by luciferase reporter assay in vitro or subcutaneous implantation and bioluminescence imaging in vivo. (G) HRE-luciferase reporter activity (RLU) in FaDu (i) and SCC1 (ii) cells under normoxia or hypoxia across four conditions: DMSO/siNTC, DMSO/siACTL6A, T-5224/siNTC, and T-5224/siACTL6A. n = 3. *** p < 0.001, **** p < 0.0001, two-way ANOVA followed by Tukey’s multiple comparisons test. (H) Representative bioluminescence images at day 21 of mice bearing subcutaneous MOC2 tumors stably expressing the HRE-luciferase reporter with shNTC (top) or shactl6a (bottom). (I) Mean radiance (photons/sec/cm2/sr) of tumors over time. n = 5-6/group. * p < 0.05, **** p < 0.0001, two-way ANOVA followed by Sidak’s multiple comparisons test. (J) Tumor volume measured over time. * p < 0.05, **** p < 0.0001. (K) Endpoint tumor weight (left) and representative image of harvested tumors (right). ** p < 0.01. All data are mean ± SEM.

Our RNA-seq data showed enrichment of the Hallmark hypoxia signature in control relative to ACTL6A-depleted cells (NES = 1.3, p = 0.043, FDR q = 0.22) (Fig. 3B) and the majority of Buffa hypoxia genes (27) were down-regulated following ACTL6A knockdown (Fig. 3C). Because the hypoxia-inducible factors HIF1α and HIF2α are the principal effectors of this program and are regulated post-translationally, we measured HIF protein levels in FaDu and SCC1 cells after transient ACTL6A knockdown under normoxia or 0.2% hypoxia. ACTL6A depletion downregulated hypoxic induction of HIF2α in both cell lines and of HIF1α in FaDu cells (Fig. 3D), which was consistent with ACTL6A occupancy of the EPAS1/HIF2α enhancer that we previously identified. Functionally, ACTL6A depletion significantly abrogated cell growth under 0.2% hypoxia in both FaDu and SCC1 (Fig. 3E).

To test the interaction ACTL6A and AP-1 transcription factor activity in regulating hypoxic adaptation, we used a hypoxia-response-element (HRE)-driven luciferase reporter (3× HRE elements upstream of a minimal promoter) and measured reporter activity following ACTL6A depletion and/or treatment with the AP-1 inhibitor T-5224 under normoxia or hypoxia (Fig. 3F). We first confirmed that T-5224 inhibits AP-1 activity in our cells, as it dose-dependently suppressed PMA-induced expression of the AP-1 early target gene FOSL1 in both SCC1 and FaDu (fig. S3A). HRE-luciferase activity was strongly induced by hypoxia in control (DMSO/siNTC) FaDu and SCC1 cells and was significantly reduced by either ACTL6A knockdown or T-5224 alone, with the combination producing the lowest reporter signal (Fig. 3G). These results support a model in which ACTL6A enhances the hypoxic response through AP-1 transcription factor activity.

Based on results of ACTL6A coordinating hypoxic adaptation in HNSCC cells and within solid tumors in TCGA, we hypothesized whether its expression directly regulates hypoxic responses in tumors *in vivo*. MOC2 cells, which are derived from a carcinogen-induced mouse oral cavity carcinoma, were stably transduced with the HRE-luciferase reporter and either control (shNTC) or ACTL6A-targeting (shACTL6A) shRNA. These were injected into flanks to generate tumors that were monitored by bioluminescence imaging (fig. S3B). At day 21, shNTC tumors displayed significantly stronger reporter signal than shACTL6A tumors (Fig. 3H), and longitudinal quantification showed significantly higher mean radiance in control tumors from day 14 onward (*p* < 0.05 at day 14, *p* < 0.0001 at day 21) (Fig. 3I) with concordant results for total photon emission (fig. S3C). This reduction in HIF reporter activity was accompanied by impaired tumor growth, with shACTL6A tumors significantly smaller by tumor volume (*p* < 0.05 at day 14, *p* < 0.0001 at days 17 and 21) (Fig. 3J) and by endpoint tumor weight (*p* < 0.01) (Fig. 3K). Together, these results establish that ACTL6A is required for HIF-dependent transcriptional activity and hypoxic tumor growth *in vivo*.

### ACTL6A depletion potentiates mitochondrial mass and function

There are no prior studies on how ACTL6A levels influence mitochondria and single cell metabolic properties of cancer cells. We initially assessed mitochondrial mass by transducing SCC1 and FaDu cells with the MitoDsRed plasmid then transiently knocked down ACTL6A. We observed significantly increased intensity of MitoDsred after ACTL6A depletion in both cell lines, indicative of enhanced mitochondrial density and mass (Fig. 4A). We then used Mitotracker deep red (MTDR) staining to study alterations in mitochondrial potential and activity in SCC1 and FaDu cells and found significantly increased MTDR staining after ACTL6A depletion (Fig. 4B). Mitochondria are the main source of reactive oxygen species (ROS), which are generally cytotoxic to cancer cells and regulated through oncogenic redox homeostasis pathways (28). We tested whether increased mitochondrial activity in cells with ACTL6A knockdown could lead to increased superoxide generation using MitoSOX red staining followed by flow cytometry. We observed a significant increase in MitoSOX intensity after ACTL6A knockdown in both SCC1 and FaDu cells (Fig. 4C).

**Figure 4.**
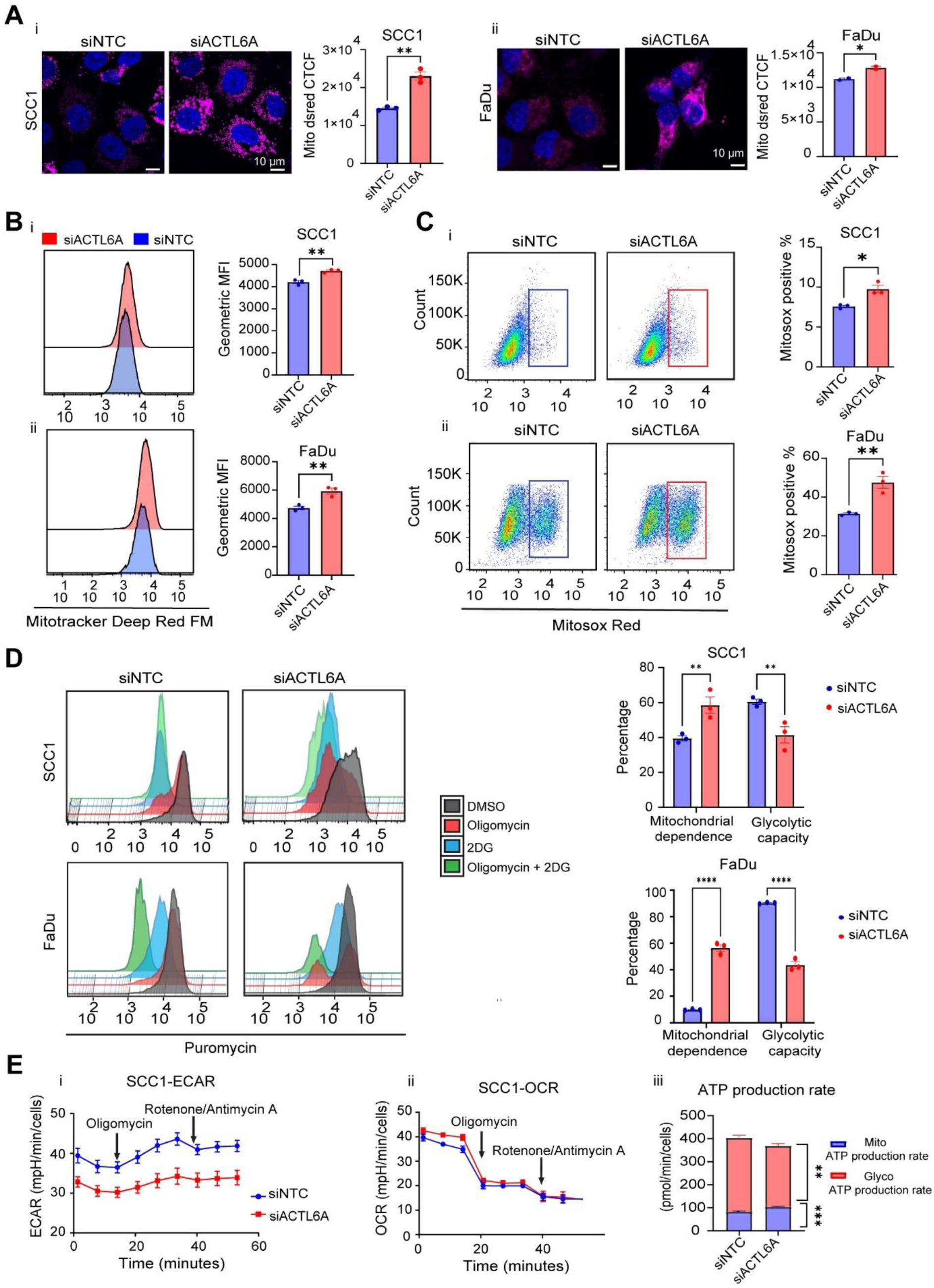
Mitochondrial function is enhanced by ACTL6A depletion. (A) Representative confocal images and quantitation of MitoDsRed (magenta) normalized to the number of nuclei stained with DAPI (blue) in SCC1 (i) and FaDu (ii) cells following transient ACTL6A knockdown. (n=3 in SCC1 and n=2 in FaDu). Data are mean ± SEM, * p < 0.05, ** p < 0.01, unpaired t tests. Scale bar: 10µm. (B) Representative histogram and quantitation of Geometric Mean Fluorescence (GMF) of Mitotracker Deep Red FM in SCC1 (i) and FaDu (ii) analyzed by flowcytometry after 72 hours transient ACTL6A knockdown. (n=3), Data are mean ± SEM, ** p < 0.01, unpaired t test. (C) Representative pseudo color graphs from flowcytometry analysis and quantitation of percentage of Mitosox positive in SCC1 (i) and FaDu (ii) cells following transient ACTL6A knockdown. N=3. Data are mean ± SEM, * p < 0.05, ** p < 0.01, unpaired t tests. (D) Histogram plots of puromycin fluorescence intensity (i) and quantitation (ii) of percentage of mitochondrial dependence and glycolytic capacity from SCENITH assay (flowcytometry based) in SCC1 (upper panel) and FaDu cells (lower panel) following ACTL6A knockdown. N=3. Data are mean ± SEM, (E) Representative extracellular acidification rate (ECAR) (i), oxygen consumption rate (ii), ATP production rate (iii) from SCC1 after transient ACTL6A knockdown using Seahorse XF Real-Time ATP Rate assay. N=2. ** p < 0.01, *** p <0.0001, two-way ANOVA followed by Sidak’s multiple comparisons test.

### ACTL6A expression alters HNSCC cell metabolism

To characterize how ACTL6A influences HNSCC cell metabolism, we performed a flow cytometry-based assay called SCENITH (single-cell energetic metabolism by profiling translation inhibition) that uses protein translation as a correlate of energy consumption within cells (29). We treated the cells with glycolysis or oxidative phosphorylation inhibitors including 2-deoxy-d-glucose (2DG) and oligomycin, respectively, or both to block total ATP production. Next, we incubated the cells with puromycin and stained with anti-puromycin antibody to assess changes in protein synthesis in SCC1 and FaDu cells ACTL6A depletion (Fig. 4D). These data showed that upon addition of oligomycin (Fig 4Di, red), MFI (mean Fluorescence intensity) of puromycin decreased more significantly with ACTL6A depletion compared with control, which is indicative of higher mitochondrial dependency and lower glycolytic capacity in siACTL6A cells (Fig. 4Dii).

Additionally, we performed a Seahorse XF Real-Time ATP Rate assay in SCC1 cells following transient ACTL6A knockdown to estimate ATP production from mitochondria and glycolysis by measuring extracellular acidification (ECAR) and oxygen consumption rate (OCR) in media upon sequential treatment with oligomycin (Complex V inhibitor) and rotenone/antimycin A (complex I and III inhibitors). The ATP rate assay revealed diversion of glycolysis to oxidative phosphorylation while total ATP production did not change with ACTL6A knockdown (Fig. 4E), which was consistent with SCENITH assay. These findings indicate that high ACTL6A levels up-regulate glycolysis and down-regulate oxidative phosphorylation, consistent with decreasing mitochondrial dependency.

ACTL6A knockdown was previously shown to down-regulate cell proliferation and spheroid formation in HNSCC (11). We examined effects of ACTL6A depletion on additional oncogenic cell phenotypes associated with cell metabolism, including cell motility (30) and cisplatin resistance (31). Using a transwell migration assay, we found that ACTL6A depletion significantly reduced cell migration in two independent HPV-negative cell lines, SCC1 and FaDu (fig. S4A). We then measured the IC50 of cisplatin in FaDu and UM-SCC2 cells, an additional HPV-negative cell line, and found ACTL6A depletion significantly increased sensitivity to cisplatin compared to control cells (fig. S4B). These findings are consistent with down-regulation of the epithelial mesenchymal transition (p < 0.001) and xenobiotic metabolism (p < 0.05) pathways observed in our GSEA (Fig. 1D).

### ACTL6A expression potentiates upstream MAPK signaling

Mitogen-activated protein kinase (MAPK) signaling pathway is a tightly regulated signaling cascade that induces AP-1 TF binding (32). By integrated analysis of RNA-seq, ATAC-seq, and ACTL6A CUT&RUN, we show that ACTL6A directly regulates chromatin accessibility at AP-1 TF sites, particularly at glycolysis and hypoxia genes. Further analysis using RNA-seq also revealed downregulation of the majority of late p-ERK responsive genes (33) in SCC1 cells with ACTL6A depletion (fig. S5A). Additionally, using our DepMap set of HPV-HNSCC cells lines, we found significant correlations between *ACTL6A* and *KRAS* (p<0.001), *NRAS* (p<0.001), and *HRAS* expression (p<0.05) (Fig. 5A). We also identified LGALS1/Galectin-1 among the top five most downregulated genes with ACTL6A knockdown (Log2FC = -1.6, Adj p-value = 8.32E-09) (fig. S5B). Galectin-1 stabilizes RAS protein anchorage to the cell membrane, which potentiates downstream signaling through MEK-ERK (34). We identified a TEAD1 binding site in the *LGALS1* promoter, which had significantly decreased accessibility in siACTL6A samples (fig. S5B). Based on this evidence, we hypothesized that ACTL6A expression also coordinates MAPK signaling to further drive AP-1 mediated gene expression. We quantified protein expression of Galectin-1, RAS, and p-ERK1/2 in FaDu and SCC1 cells following ACTL6A knockdown, which showed reduced expression of these proteins after siACTL6A treatment (Fig. 5B). We also confirmed that ACTL6A depletion down-regulates *Ras* gene expression in SCC1 cells (Fig. 5C).

**Figure 5.**
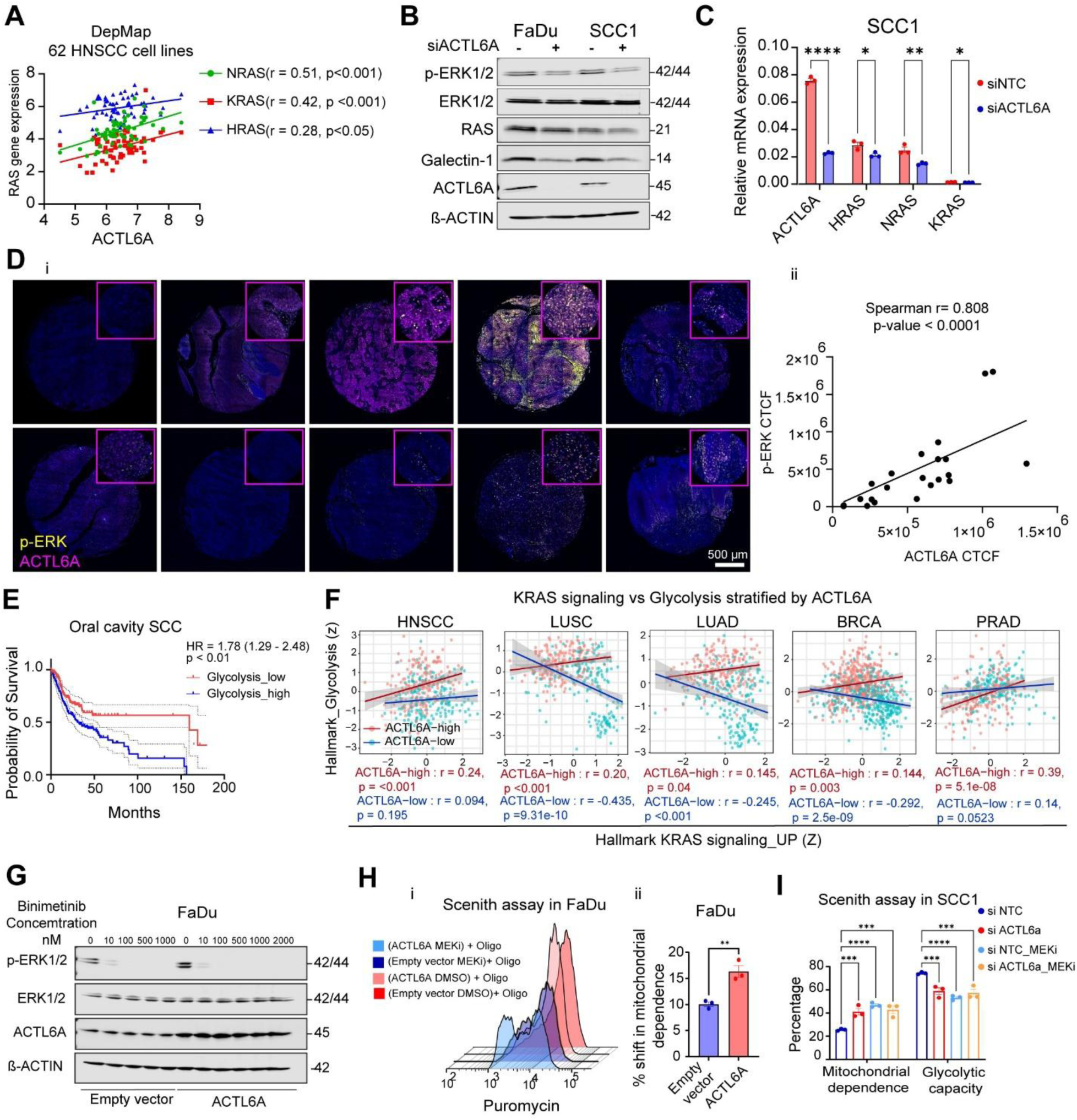
ACTL6A potentiates ERK phosphorylation and sensitizes HNSCC cells to the metabolic effects of MEK inhibitors. (A) Correlation analysis between ACTL6A and RAS genes in 62 HNSCC cell lines from DepMap. (B) Representative Western blot images of β-actin, ACTL6A, Galectin-1, RAS, total ERK1/2, and p-ERK1/2 in FaDu and SCC1 cells following transient ACTL6A knockdown. (C) qRT-PCR analysis individual Ras genes following ACTL6A depletion or non-targeting siRNA in SCC1 cells. (D) Representative confocal images (i) and non-parametric spearman correlation analysis (ii) of corrected total cell fluorescence for ACTL6A (magenta) and p-ERK (yellow). Immunofluorescence staining was done on paraffin embedded freshly cut tissue array sections from either HNSCC or normal tongue tissue (N=24 cores) (E) Glycolysis enrichment scores were generated using GSVA and used to stratify HNSCC patients in TCGA by median enrichment. Kaplan-Meier curve showing decreased overall survival in patients with higher glycolysis compared to lower glycolysis. (F) Correlation between Hallmark KRAS signaling (up) and Hallmark glycolysis enrichment scores (z-scores) across five TCGA cohorts (HNSCC, LUSC, LUAD, BRCA, and PRAD), with tumors stratified into ACTL6A-high (red) and ACTL6A-low (blue) groups. Correlation coefficients (r) and p values are shown for each group; shaded bands indicate the 95% confidence interval of the linear fit. (G) Representative Western blot of p-ERK1/2, total ERK1/2, ACTL6A, and β-actin in FaDu cells expressing empty vector or ACTL6A following 2-hour treatment with the indicated concentrations of Binimetinib. (H) Representative histograms of puromycin fluorescence intensity (i) and quantitation (ii) of the percentage shift in mitochondrial dependence from the SCENITH assay in FaDu cells expressing empty vector or ACTL6A treated with DMSO or MEKi. n = 3, ** p < 0.01, unpaired t test. (I) Geometric mean fluorescence intensity of puromycin was measured in different conditions to calculate mitochondrial dependence and glycolytic capacity in SCC1 cells following ACTL6A depletion and 2-hour treatment with either DMSO or 10 nM Binimetinib (MEK inhibitor). n = 3, *** p < 0.001, **** p < 0.0001, two-way ANOVA followed by Tukey’s multiple comparisons test. Data are mean ± SEM

To test if ACTL6A is correlated with p-ERK expression in human HNSCC tissue, we performed immunofluorescence staining for both proteins on freshly cut tissue microarray (TMA) of ten HNSCC tumors and two normal upper aerodigestive tract mucosa specimens with 2 cores per specimen (Fig. 5D,i). We found a strong, significant correlation between ACTL6A and p-ERK expression in the TMA (r = 0.808, p<0.001) (Fig. 5D,ii). These data link ACTL6A with induction of up-stream MAPK signaling, indicating coordinated regulation of AP-1 signaling through distinct mechanisms.

### ACTL6A potentiates MAPK/AP-1 signaling to program HNSCC cell metabolism

To define the clinical implications of ACTL6A expression on glycolysis, we analyzed oral cavity SCC patients in TCGA as this is the most common HPV-subsite (n = 316). Patients were stratified by median glycolysis gene set variation analysis (GSVA) enrichment scores (table S4). Patients with high glycolysis had a 78% increased risk of death compared to those with low glycolysis, defining its importance as a prognostic indicator (HR = 1.78, p<0.01) (Fig. 5E). ACTL6A expression was also significantly higher in those with high glycolysis compared to low glycolysis (p < 0.05) (fig. S5E). MAPK/ERK signaling is a driver of glycolysis in both normal and cancer cells (35) and our data has shown that ACTL6A is a key regulator of both upstream MAPK pathway activation and chromatin accessibility at AP-1 sites within glycolytic enzymes. Therefore, we hypothesized that ACTL6A expression regulates the relationship between MAPK signaling and glycolytic metabolism in human tumors. We generated Hallmark KRAS signaling (up) and Hallmark glycolysis enrichment scores for each patient and correlated them within ACTL6A-high and ACTL6A-low tumors across five TCGA solid tumor cohorts. In ACTL6A-high tumors, KRAS signaling was significantly and positively correlated with glycolysis in every cohort (HNSCC *r* = 0.24, *p* < 0.001; LUSC *r* = 0.20, *p* < 0.001; LUAD *r* = 0.145, *p* = 0.04; BRCA *r* = 0.144, *p* = 0.003; PRAD *r* = 0.39, *p* = 5.1E-08). In contrast, this relationship was absent or significantly inverted in ACTL6A-low tumors (HNSCC *r* = 0.094, *p* = 0.195; LUSC *r* = −0.435, *p* = 9.31E-10; LUAD *r* = −0.245, *p* < 0.001; BRCA *r* = −0.292, *p* = 2.5E-09; PRAD *r* = 0.14, *p* = 0.0523) (Fig. 5F).

Based on these correlations, we hypothesized that ACTL6A’s metabolic effects are mediated through MAPK signaling and may be susceptible to MEK inhibitors. To test this, we first treated either FaDu control or ACTL6A overexpressing cells with either DMSO or 10nM Binimetinib, a small molecule inhibitor of MEK1/2, (hereafter as MEKi) for 2 hours. We selected this concentration based on the ability of MEKi in inhibiting ERK phosphorylation (Fig. 5G). We then performed SCENITH assay, which showed ACTL6A overexpressing cells had a significantly larger shift toward mitochondrial dependence compared to control cells (Fig. 5H), indicating greater sensitivity to the metabolic effects of MEKi. We then compared the effects of MEKi on ACTL6A depleted and control SCC1 cells. Using SCENITH, we found that control cells had increased mitochondrial dependency and decreased glycolytic capacity with MEKi treatment (p<0.001). In contrast, ACTL6A-depleted cells had mitochondrial dependency and glycolytic capacity that was similar to control cells after MEKi treatment and had no change in metabolism with MEKi treatment (Fig. 5I), suggesting MAPK/ERK-induced effects on potentiation glycolysis were absent after ACTL6A depletion. These data show that ACTL6A expression is necessary for the metabolic effects of MEK/ERK.

### An ACTL6A-dependent transcriptional program marks hypoxic, spatially isolated niches in human HNSCC

To determine whether an ACTL6A regulates intratumoral hypoxia and metabolism, we applied the CosMx 1000-plex single-cell spatial transcriptomics platform to 7 tumors from patients with oral cavity and larynx HPV (-) HNSCC. This dataset encompassed 70,736 malignant cells with annotated immune, stromal, and endothelial cell subtypes. Because ACTL6A is not represented within the gene panel and differences in expression may not fully account for transcriptional output, we generated an ACTL6A-dependent gene signature based on the 265 genes with decreased expression and chromatin accessibility on RNA-seq and ATAC-seq, respectively, after ACTL6A depletion and direct ACTL6A occupancy on CUT&RUN (Fig. 2D). Of these genes, 29 were present in the CosMx panel (Table S4). Per-cell ACTL6A activity was calculating using a scaled-mean score on SCT-normalized expression data in malignant cells. Quality control was performed by comparing results to UCell scoring (36) (fig. S6A,B) and there were no differences in transcript variation based on ACTL6A activity score (fig. S6C,D). We used a similar approach to score per-cell hypoxia and MAPK signaling pathways using the Hallmark Hypoxia and KRAS Up pathway gene sets after excluding genes overlapping with the ACTL6A activity gene set (4 of 38 and 3 of 52, respectively, Table S4). ACTL6A activity was non-uniform across tumors and strongly co-localized with regions of high Hypoxia scores (Fig. 6A). Malignant cells with highest quartile ACTL6A activity scores were also enriched in genes associated with basal progenitor cells, including TACSTD2, CD44, and ITGB4 (fig. S7A).

**Figure 6.**
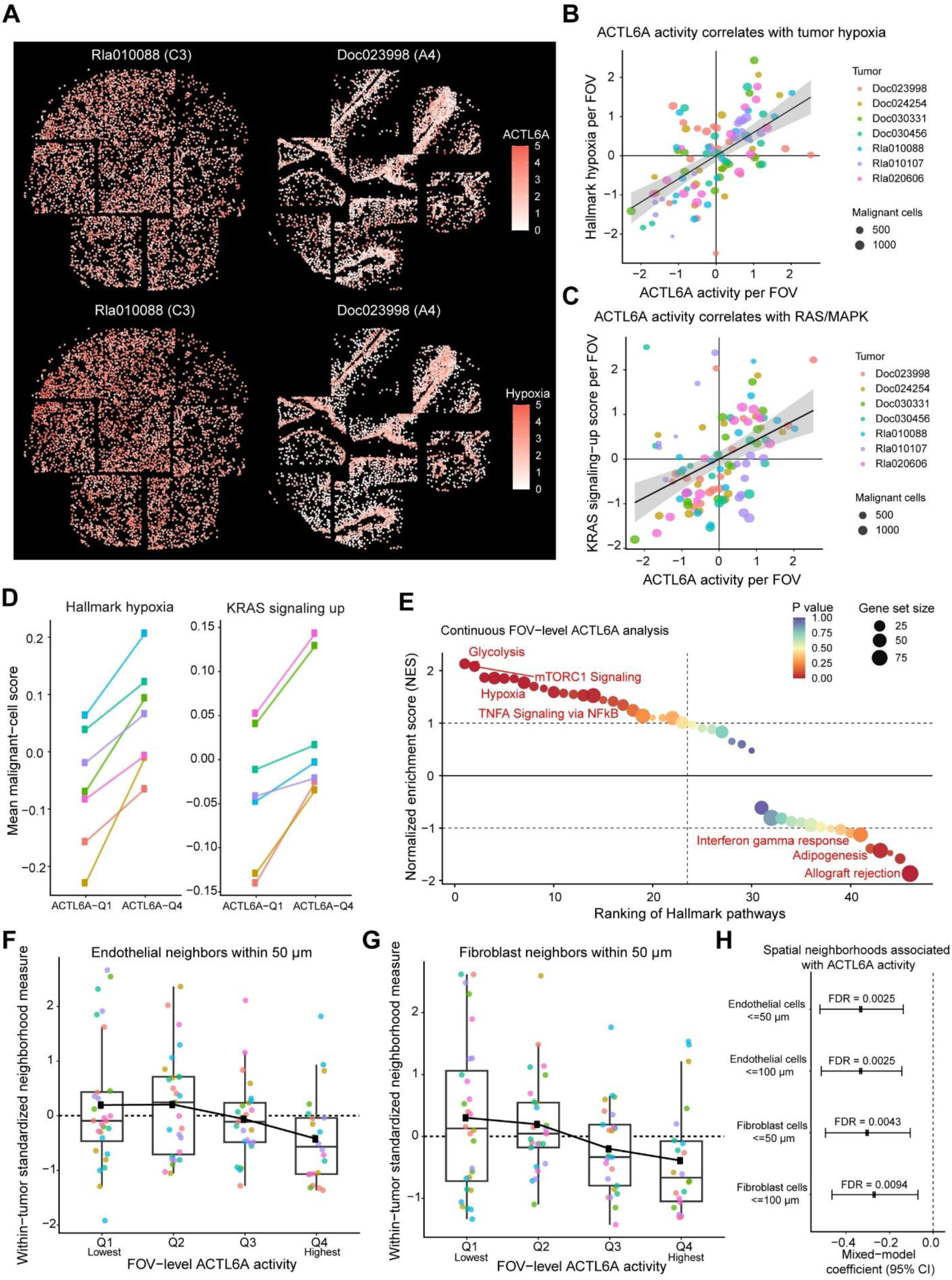
An ACTL6A-dependent transcriptional program marks hypoxic, vessel poor niches in HNSCC. (A) Representative cores from CosMx 1000-plex spatial transcriptomics analysis indicating per-cell ACTL6A activity and Hypoxia program scores in tumors with high (Rla010088, core C3) and low (Doc023998, core A4) mean ACTL6A activity. FOV-level correlations between ACTL6A activity and (B) Hallmark hypoxia and (C) Hallmark KRAS signaling-up programs. Each point represents one FOV (n = 102 FOVs) with a mean program score across malignant cells within each FOV. Regression lines show linear fits with 95% confidence intervals. (D) Mean Hallmark hypoxia (left) and KRAS signaling-up (right) scores in malignant cells in the highest (Q4) versus lowest (Q1) quartile of ACTL6A activity, defined separately within each tumor. Each line connects paired values from a single tumor (n = 7). (E) Hallmark pathway rank plot ordered by normalized enrichment score (NES). FOV-level (F) endothelial and (G) fibroblast neighbor counts within 50 µm of malignant cells, stratified by within-tumor quartile of ACTL6A activity. Boxplots show FOV-level distributions (center line, median; box, interquartile range; whiskers, 1.5 × IQR); individual points denote FOVs, colored by tumor. (H) Standardized coefficients (95% CI) from linear mixed-effects models of FOV-level endothelial or fibroblast neighbor abundance on continuous ACTL6A activity, with random intercepts for tissue core. Benjamini–Hochberg FDR calculated across the four models.

ACTL6A activity scores were correlated with Hypoxia and KRAS scores using field of view (FOV) averaged scores (102 FOVs, >13 FOVs per tumor). Using a mixed-effects models to account for tumor and tissue-core structure, FOV-level ACTL6A activity was strongly associated with increasing Hypoxia score (β = 0.592, FDR = 1.62x10^-10^) and increasing KRAS score (β = 0.357, FDR = 3.03x10^-4^) (Fig. 6B,C). These results retained significance at the tumor level (fig. S7B). We then compared malignant cells in the highest and lowest quartiles of ACTL6A activity within each tumor. ACTL6A-high cells exhibited significantly higher mean Hypoxia and KRAS scores in all seven tumors (p=0.0156) (Fig. 6D), indicating ACTL6A transcriptional activity may broadly generalize across HNSCCs.

We then performed an FOV-level GSEA using the Hallmark gene set while modeling ACTL6A activity as continuous variable. Genes within the ACTL6A activity gene set were excluded from the Hallmark pathway gene sets. We identified significant positive enrichment of multiple metabolic and stress-response pathways in regions with higher ACTL6A activity. We identified strong positive associations for Glycolysis (NES = 2.13, FDR = 0.008) and Hypoxia (NES = 1.77, FDR = 0.0168) (Fig. 6E).

Given strong associations between ACTL6A expression and hypoxia, we hypothesized that increased ACTL6A activity scores may allow cells to persist further from endothelium and stroma and drive spatial isolation. The number of endothelial cells and fibroblasts within 50- and 100-µm radii was calculated for each malignant cell. These were averaged across FOVs, which were divided by ACTL6A activity quartiles. ACTL6A activity was associated with fewer endothelial neighbors within both 50 µm (β = −0.309, FDR = 0.0025) and 100 µm (β = −0.328, FDR = 0.00076) as well as fibroblast abundance within 50 µm (β = −0.300, FDR = 0.0043) and within 100 µm (β = −0.278, FDR = 0.0094) (Fig. 6F-H). These spatial relationships were demonstrated across 5 of 7 tumors (fig. S7C).

### Targeting treatment resistant HNSCC tumors with ACTL6A depletion and oxidative phosphorylation inhibitors

Our epigenomic and functional data indicate that ACTL6A is a key driver of glycolysis and hypoxic plasticity in HNSCC cells. We used TCGA to calculate ratios of glycolysis gene set to oxidative phosphorylation gene set expression across solid tumors and found significant positive correlations with ACTL6A expression in 11 of 24 tumor datasets, including HNSCC (rho = 0.17, FDR = 1.5x10^-4^), BLCA (rho = 0.15, FDR = 3x10^-3^), BRCA (rho = 0.14, FDR = 1.1x10^-5^), KIRP (rho = 0.32, FDR = 5.2x10^-6^), LIHC (rho = 0.56, FDR < 2.2x10^-16^), and PAAD (rho = 0.4, FDR < 2.2x10^-16^) (Fig. 7A, fig. S8C). Notably, ACTL6A depletion induces significant up-regulation of mitochondrial mass and function, suggesting these may be compensatory mechanisms that overcome decreased glycolytic output and that these cells may be dependent on mitochondrial sources of energy production. We therefore hypothesized that decreasing ACTL6A levels could sensitize cells to mitochondrial inhibitors. We used IACS-010759, a potent inhibitor of oxidative phosphorylation that targets mitochondrial complex I and has been in clinical trials for solid tumors (37) to assess chemosensitivity of HNSCC cell lines to mitochondrial inhibitors. We assessed IC50 to IACS-010759 after ACTL6A knockdown in FaDu cells, which showed significantly enhanced sensitivity to low doses of drug at 25 and 50 nM in siACTL6A cells (Fig. 7Bi). Conversely, FaDu cells over-expressing ACTL6A were significantly more treatment resistant to IACS-010759 relative to control cells, even at high drug concentrations of 500 nM (Fig. 7Bii). We then assessed sensitivity of MOC2, which are highly proliferative and resistant to cisplatin, radiation, immunotherapy, and combination therapies (38,39). We generated a stable ACTL6A knockdown using shACTL6A with a ∼50% reduction in ACTL6A levels (fig. S8C). We found shACTL6A MOC2 cells had significantly increased sensitivity to IACS-010759 relative to control cells across a range of drug concentrations (Fig. 7Biii).

**Figure 7.**
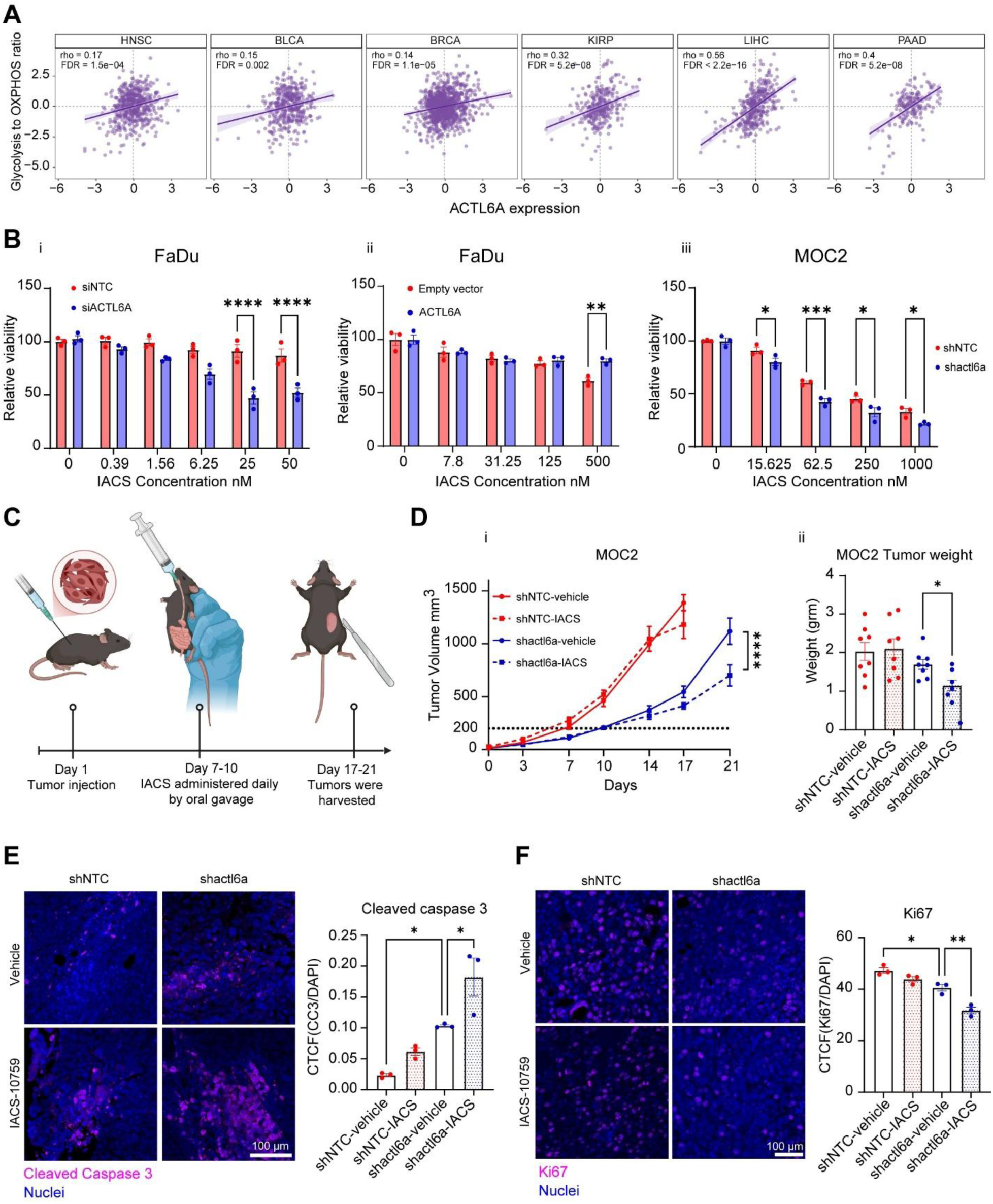
ACTL6A knockdown sensitizes HNSCC cells to treatment with the complex I inhibitor IACS-010759. (A) Correlation between Glycolysis to Oxidative phosphorylation gene set expression and ACTL6A expression across six TCGA cohorts (HNSCC, BLCA, BRCA, KIRP, LIHC, PAAD). Spearman r and FDR-corrected p values are shown for each group. (B) Relative viability of FaDu cells following ACTL6A transient knockdown (i), or ACTL6A overexpression (ii), and (iii) MOC2 cells following shACTL6A was measured using SRB assay after incubation with either DMSO or indicated concentrations of IACS-010759 for 5 days. N=3. * p < 0.05, ** p < 0.01, *** p <0.0001, **** p <0.0001, two-way ANOVA followed by Sidak’s multiple comparisons test. (C) Schematic showing the timeline for injection of tumor cells and administration of IACS-010759. (D) (i) Tumor volume was measured at indicated time points after injection with MOC2 cells expressing shACTL6A or control vector. When tumor size reached 200 mm^3^ (dashed line), mice were treated with either control vehicle or IACS-010759 by oral gavage. N=9-10/group, ns p > 0.05, ****p < 0.0001, two-way ANOVA followed by Sidak’s multiple comparisons test. (ii) Tumor weight was measured at study endpoint. ns p > 0.05, * p < 0.05, unpaired t tests. (E) Representative confocal images from tumor tissue stained for cleaved caspase 3 (cc3) (magenta) and DAPI (blue) (left) and quantification (right) of corrected total cell fluorescence of cc3 normalized to DAPI in indicated samples. N=3. * p < 0.05, one-way ANOVA followed by Tukey’s multiple comparisons test. Scale bar: 100µm. (F) Representative confocal images from mice tissue stained for Ki67 (magenta) and DAPI (blue) (left) and quantification (right) of corrected total cell fluorescence of Ki67 normalized to DAPI in indicated samples. N=3. * p < 0.05, ** p < 0.01, one-way ANOVA followed by Tukey’s multiple comparisons test. Scale bar: 100µm.

Based on the synergistic effects of ACTL6A knockdown and IACS-010759 treatment *in vitro*, we investigated the effect of this combined treatment on tumor growth *in vivo.* We performed a heterotopic tumor model using control (shNTC) or ACTL6A-depleted (shACTL6A) MOC2 cells subcutaneously injected into B6J.Rag2 Il2rg double KO mice. Mice were treated with either vehicle or 10 mg/kg IACS-010759 by oral gavage starting when average tumor size reached 200 mm^3^ (Fig. 7C). After treatment onset, there was a significant reduction in tumor size with IACS-010759 treatment compared to vehicle in ACTL6A-depleted tumors while no change in tumor volume was observed in control tumors (Fig. 7Di). Tumors were harvested upon reaching 1000 mm^3^ in the control group and endpoint measurement of tumor weight revealed similar results, with IACS-010759 treatment leading to significantly lower tumor weight only in ACTL6A-depleted tumors (Fig. 7Dii). Immunofluorescence staining showed both a significant increase in apoptosis measured via cleaved caspase 3 (CC3) and reduced proliferation measured via Ki67 expression with IACS-010759 treatment only in the ACTL6A-depleted tumors (Fig. 7E,F). These data indicate targeting ACTL6A in combination with inhibitors of oxidative phosphorylation or mitochondrial function may be a treatment strategy for treatment resistant HNSCC.

## DISCUSSION

Uncovering pathways that regulate cancer cell metabolism provides insights into oncogenesis and offers therapeutic targets. As cancer cells shift metabolism from oxidative phosphorylation to aerobic glycolysis, they reduce their exposure to ROS and increase glucose turnover, both of which enhance proliferation (3). We have discovered that expression of ACTL6A, a gene on the 3q26 amplicon over-expressed in several cancers, acts as a switch that suppresses mitochondrial dependency and up-regulates aerobic glycolysis, leading to decreased ROS, increased hypoxic adaptation, and increased growth within spatially isolated tumor niches. These effects are mediated by coordinated mechanisms that amplify ERK signaling through up-regulation of Ras and Galectin-1 as well as maintaining chromatin accessibility at AP-1 transcription factor binding sites. This pathway can be modulated for therapeutic benefit, either through MEK inhibition or a synthetic lethal strategy targeting ACTL6A expression with inhibitors of oxidative phosphorylation. We also report the first study of ACTL6A chromatin occupancy using CUT&RUN, offering insights into its SWI/SNF dependent and independent genomic regulation. These data establish ACTL6A as a key regulator of the Warburg effect and hypoxia in HNSCC and propose ACTL6A inhibition, either through SWI/SNF inhibition or development of a direct inhibitor, as a potential treatment strategy for HNSCC.

The RAS/RAF/MAPK pathway has been linked to activation of glycolysis in several cancers. Notably, KRAS over-expressing models of pancreatic cancer significantly up-regulate glucose uptake and glycolysis gene expression (40). Similarly, MEK inhibitors have been shown to significantly decrease glycolysis in a range of tumors (41). Up-regulation of oxidative phosphorylation and mitochondrial biogenesis are key mechanisms of resistance to MAPK pathway inhibitors, which can be overcome with administration of metabolic inhibitors to rescue their anti-tumor effects (42–44). Our study demonstrates a novel mechanism through which cancer cells up-regulate aerobic glycolysis by demonstrating ACTL6A directly occupies AP-1 transcription factor binding sites and enhances chromatin accessibility at glycolysis gene promoters. ACTL6A expression levels mediate access to these sites as well as regulating phosphorylation of ERK, thereby enhancing both AP-1 transcription factor activity and chromatin access. MEK inhibitors also showed a role in increased mitochondrial dependency to a much stronger degree in ACTL6A over-expressing tumors, suggesting their potential for treating a large subset of HNSCC tumors with increased ACTL6A expression. As many RAS/RAF/MAPK inhibitors are clinically approved for use in other cancers (45), it raises the possibility of clinical trials in HNSCC after stratifying tumors using hypoxia imaging or ACTL6A expression (46,47).

ACTL6A-mediated up-regulation of ERK signaling and chromatin accessibility of AP-1 binding sites has not previously been reported. While our ATAC-seq data recapitulated increased TEAD/YAP chromatin accessibility with increasing ACTL6A expression levels (10,11), AP-1 signaling is likely to be more important for maintenance of the metabolic phenotype in our study. YAP is known to promote cell proliferation and oncogenesis in HNSCC (48), but its activation is generally modulated by cellular energy states rather than having a direct impact on them (49). For instance, several studies have shown that high glucose turnover and aerobic glycolysis activate YAP while energy stress leads to its inhibition (50–52). There are relatively few studies linking SWI/SNF and MAPK pathways, but two recent papers support SWI/SNF potentiating MAPK signaling. Notably, Wolf et al (2023) showed that multiple SWI/SNF subunits are enriched at AP-1 transcription factor binding sites and cooperate to maintain chromatin accessibility (53). Miguel et al (2023) showed that resistance to tyrosine kinase inhibitors in lung cancer was mediated by up-regulation of SWI/SNF chromatin binding at downstream MAPK genes, which could be overcome using inhibition of the SWI/SNF subunits SMARCA2 and SMARCA4 (54). We now provide evidence that ACTL6A, which is over-expressed in a variety of cancers, can also regulate accessibility of AP-1 sites in addition to potentiating MAPK signaling. Additionally, our co-IP experiment performed in native cells with no protein tagging showed a strong association between ACTL6A and c-Fos, which indicates coordination between ACTL6A and AP-1 dependent transcription factors within chromatin. ACTL6A is also a component of other complexes, including the histone acetyltransferase complex TIP60 and chromatin remodeling complex INO80 (25,55) and follow-up studies are needed to demonstrate the extent to which SWI/SNF, TIP60, INO80 coordinate oncogenic phenotypes in ACTL6A amplified HNSCC.

Modifying ACTL6A expression had significant effects on mitochondrial dependency in HNSCC cells, with low levels of ACTL6A abrogating hypoxic cell growth as well as sensitizing cells to cytotoxicity using the oxidative phosphorylation inhibitor IACS-010759. Increased mitochondrial dependence after ACTL6A knockdown is likely mediated by inability to up-regulate glycolysis in HNSCC cells rather than a change in mitochondrial function. Both MAPK signaling and SWI/SNF have been linked to changes in mitochondrial metabolism in other cancers. In melanoma, BRAF inhibition was shown to up-regulate oxidative phosphorylation through increased PGC1α and MITF expression (56). SMARCA4 deficient lung cancers were also shown to up-regulate oxidative phosphorylation through activation of PGC1α (57); however, PGC1α and MITF did not have detectable expression in SCC1 cells even after ACTL6A knockdown (see RNA-seq dataset), suggesting other transcriptional mediators may be at play. Finally, on our Seahorse analysis, there was a much greater decrease in basal ECAR than increase OCR in SCC1 cells with ACTL6A knockdown, suggesting ACTL6A levels primarily impact glycolysis. The extent to which ACTL6A acts directly on mitochondria needs further exploration and will be the subject of follow-up studies.

IACS-010759 was shown to significantly suppress growth of the treatment resistant MOC2 cell line after ACTL6A knockdown. This is similar to its reported efficacy for SMARCA4 mutated lung cancers reported by Deribe et al. (2018) (57), suggesting oxidative phosphorylation inhibitors generally have potential for treatment of SWI/SNF depleted tumors. IACS-010759 has undergone phase I clinical trials in a wide range of solid tumors and acute myeloid leukemia (AML) with some reports of efficacy but significant dose limiting toxicities, including lactic acidosis and peripheral neuropathies (37). Notably, patients were treated without screening for mutations or baseline metabolic status of tumors. While clinical development of IACS-010759 is limited by toxicity, the results in this study raise the question of whether SWI/SNF deficient tumors may be more susceptible to its effect in a lower therapeutic window where adverse effects are more tolerable. Additionally, inhibitors of other components of aerobic metabolism may also be studied. CPI-613 is a tricarboxylic acid (TCA) cycle inhibitor that has been safely delivered to humans in a clinical trial for pancreatic adenocarcinoma (58). Gamintrinib is a mitochondrial matrix inhibitor with potentially improved toxicity profile compared to IACS-010759 under clinical trial for solid tumors (59). The extent to which SWI/SNF deficient malignancies respond to metabolic inhibitors will be the subject of further study.

In conclusion, we have identified ACTL6A, a SWI/SNF subunit over-expressed in several cancers, as a key mediator of hypoxia and the Warburg effect, allowing malignant cells to persist in spatially isolated niches that predispose to tumors to treatment resistance. ACTL6A levels act as an epigenetic switch that generates a chromatin structure permissive for metabolic plasticity and depleting ACTL6A induces a state of mitochondrial dependence and sensitizes cells to oxidative phosphorylation inhibitors. These effects are induced by coordinated up-regulation of AP-1 signaling through induction of MAPK signaling as well as maintaining chromatin access at AP-1 transcription factor binding sites at glycolysis and hypoxia gene enhancers. These results uncover a basic chromatin architecture underlying cell metabolism and establish a therapeutic pathway for targeting SWI/SNF amplified malignancies.

## List of Supplementary Materials

Supplementary Methods

Fig. S1 to S8

Table S1 to S4

## Acknowledgments

We would like to thank Dr. Gerald Crabtree for providing ACTL6A over-expressing plasmids and additional reagents. We would like to thank Dr. Daria Gaykalova for technical insights and advice on the manuscript.

## Funding

National Institutes of Health grant R03DE034489 (AF)

National Institutes of Health grant R35DE030054 (JS)

American Cancer Society grant IRG-23-1074369-01-IRG (AF)

VA Palo Alto Early Career Development Award (AF)

Stanford Cancer Institute (AF, JS)

## Author contributions

AF, MM, and JS were responsible for conceptualization of the project and supervision of the experiments. AF, MM, ARS, IAM, SM, KL and MZ performed all experiments, including *in vitro*, *in vivo*, and sequencing. MM, KL, ARS, IAM, SM, MZ, JCW, KFC, and AF were responsible for data analysis. AF and MM wrote the original manuscript. JCW, KFC, and JS provided critical analysis of the manuscript for editing.

## Data and materials availability

The ATAC-seq, RNA-seq, and CUT&RUN sequencing datasets generated for SCC1 cells have been deposited in the GEO database under accession code GSE305164. Source data related to the experiments and the Seurat file used for spatial transcriptomics analysis have been deposited to the Dryad data repository using a partnership with Stanford University (DOI:10.5061/dryad.1rn8pk16b). All data are available in the main text or the supplementary materials.

## SUPPLEMENTARY MATERIALS AND METHODS

### Cell lines and culture conditions

FaDu and UM-SCC-1 cell lines were purchased from ATCC (American Type Culture Collection) and Sigma-Aldrich respectively. UM-SCC-2 cells were a gift from Dr. Thomas Carey of the University of Michigan. MOC2 (Mouse oral squamous cell carcinoma) cells were a gift from Ravi Uppaluri. All cell lines were cultured in DMEM-F12 supplemented with L-glutamine, HEPES, 10% FBS, and 100 IU/ml penicillin 100µg/ml streptomycin and incubated at 37°c in a 5% CO2 incubator. Cell line authentication was done by ATCC cell authentication service using STR profiling. Cells were tested frequently for the presence of Mycoplasma using Bulldog Bio Inc E-MYCO mycoplasma PCR kit.

### Transwell migration assay

10 µg/ml fibronectin in sterile DI water was added to transwell inserts with 6.5mm diameter and 8µm pore size and incubated at 37°c for 2 hours. FaDu and SCC1 cells were suspended in serum free media at 500000/ml density. After removing fibronectin solution from the inserts, 200 µl cell suspension was added to the upper chamber while lower chamber was filled with 600 µl complete growth media. Cells were incubated for 24 hours at 37°c in a 5% CO2 incubator. Then, medium was removed and inserts were washed gently with PBS twice and non-invaded cells were removed from the membrane by cotton tipped applicators. Next, the cells were fixed using 10% NBF (neutral buffered formalin) for 5 minutes at RT. Then the cells were washed with PBS and permeabilized with 100% ethanol for 20 minutes and stained with 0.5% crystal violet in 25% methanol in DI water for 15 minutes. Next, transwells were washed with DI water and air dried at RT overnight. Then membranes were cut and mounted on microscope slides with mounting medium. Imaging was done using ZEISS Axio Imager 2.

### ACTL6A transient Knockdown

Reverse Transient knockdown was done using ON-TARGET plus human ACTL6A SMART pool siRNAs (l-008243-00-0005, Dharmacon). Briefly, either ACTL6A or non-targeting control siRNA (ON-TARGET plus Non-targeting Pool, D-0001810-10-05, Dharmacon) was added at 100 nM concentration to 150 µl serum free OPTIMEM medium. Next, 10 µl transfection reagent (X-tremeGENE™ 360, Sigmaaldrich) was mixed with OPTIMEM medium and incubated at RT for 30 minutes. Then the mixture was added dropwise to a 6 well plate contained 1.5 ml growth OPTIMEM with 10% FBS. Next FaDu and SCC1 cells were transferred to the wells at density of 200,000 cells/well. After 72 hours the cells were collected for downstream experiments and analysis.

### ACTL6A stable knockdown

Two individual SMARTvector mouse lentiviral shRNAs (Dharmacon, V3SM11241-235868768 and V3SM11241-237525934) against mouse actl6a were used for stable actl6a knockdown in MOC2 cells. Briefly, HEK293T cells were cultured in a 100mm culture plate in DMEM-F12 supplemented with 10% FBs and 100 IU/ml penicillin 100µg/ml streptomycin. The next day, 10 µg shactl6a plasmid, 5 µg of each packaging vector (pMD2.G and psPAX2), and 50 µg PEI were mixed with 500 µl serum free OPTIMEM medium and incubated at RT for 30 minutes. Then the mixture was added to the cells dropwise. The next day, the medium was replaced with fresh complete growth medium. Viral particles containing media was collected at 48 and 72 hours, and filtered by 0.45µm PES syringe filter then concentrated using Amicon Ultra centrifugal filter at 3000 rpm for 10-20 minutes at 4°c. To transduce MOC2 cells, they were cultured in 6 well plate for 24 hours, then concentrated viral particles were added to their medium with 10 µg/ml polybrene. After 48 hours, selection was done using 2 µg/ml Puromycin.

### ACTL6A overexpression

Lentiviral vector N106 cloned with ACTL6A cDNA, a gift from Dr. Crabtree’s lab, was used for ACTL6A overexpression in FaDu cells. Lentiviral production and transduction were done as described in “ACTL6A stable knockdown” and 10 µg/ml Blasticidin was used for selection.

### Mitochondrial labeling

pLV-mitoDsRed lentiviral vector (a gift from Pantelis Tsoulfas (Addgene plasmid # 44386; http://n2t.net/addgene:44386; RRID: Addgene 44386)) was used to label mitochondria. In brief, FaDu and SCC1 were cultured in 6 well plates until about 50% confluent. Then lentiviral particles carrying pLV-mitoDsRed vector and 10 µg/ml Polybrene were added to the cells and incubated for 48 hours. Next DsRed positive cells were sorted using BD FACSAria sorter. Labeled FaDu and SCC1 cells were cultured in Thermo Scientific™ Nunc™ Lab-Tek™ II Chamber Slides followed by ACTL6A transient knockdown for 72 hours. Then the cells were washed with cold PBS twice and fixed with 10% NBF for 15 minutes. After washing with PBS for 3 times, permeabilization was done using 0.03% triton 100x in PBS for 10 minutes. The chambers were washed with PBS and mounted with prolong gold antifade mounting reagent with DAPI (Invitrogen; P36941). Imaging was done using ZEISS LSM700 confocal microscope. CTCF (corrected total cell fluorescence) for DsRed in each field was measured in Fiji 2.16.0 and normalized to the total number of cells per field.

### MitoTracker Deep Red FM and MitoSOX Red staining

MitoTracker Deep Red FM (MTDR) and MitoSOX (MedChemExpress; Cat. No.: HY-D1783 and HY-D1055 respectively) staining were done following manufacturer’s protocol. Briefly, FaDu and SCC1 cells, Following ACTL6A transient knockdown, were incubated with either 100nM MTDR or 5µM MitoSOX in Serum-free medium for 30 minutes at RT. Then, cells were washed with medium twice for 5 minutes. Flowcytometry was done using BD FACSymphony A5. FlowJo-V10 was used for analysis.

### Seahorse XF Real-Time ATP Rate Assay

Real-time cellular metabolism was analyzed using an Agilent Seahorse XFe96 analyzer (Agilent Technologies, USA) to measure oxygen consumption rate (OCR), extracellular acidification rate (ECAR), total ATP production rate, and the relative contributions of glycolytic and mitochondrial pathways. Briefly, SCC1 cells were seeded at 20,000 cells/well in a Seahorse XF96 culture microplate and maintained in their growth medium for 24 hours at 37°C under 5% CO₂. A day prior to the experiment, the Seahorse XF96 sensor cartridge was hydrated with 200 ul water and incubated in a non-CO2 incubator at 37°C. Two hours before the experiment, the water was replaced with a Seahorse XF calibrant, and the cartridge re-incubated at 37°C (non-CO2). On the day of the experiment, Seahorse XF DMEM (pH = 7.4) was supplemented with 0.5 mM pyruvate, 2.5 mM glutamine and 10mM glucose (Agilent, USA). Culture medium was replaced with 180 ul/well of supplemented Seahorse XF DMEM medium. Cells were incubated in a non-CO2 incubator at 37°C for 1 hour. During this incubation period, oligomycin and Rotenone/Antimycin were prepared in the Seahorse medium to achieve final concentration of 20 uM upon injected. The hydrated sensor cartridge was loaded with 20 ul and 22 ul of Oligomycin and rotenone/ antimycin A respectively in ports A and B and calibrated for 30 minutes. The calibration plate was replaced with the cell culture plate upon calibration completed. Measurements followed sequential injection of inhibitors using the default program. Cell numbers were quantified post-assay using SRB assay for OCR/ECAR normalization. Data analysis was done using Wave 2.6.3 software.

### SCENITH assay

SCENITH assay was performed according to the protocol outlined by Arguello et al. (2020)(29). Briefly, 72 hours after transient ACTL6A knockdown, the cells were treated with DMSO, 2µM oligomycin (O), 100 mM 2-Deoxy-D-glucose (2DG) (Medchemexpress, Cat. No: HY-13966) or combination of oligomycin and 2DG (2DGO). After 15 minutes incubation, Puromycin was added to the cells at final concentration of 10µg/ml and incubation continued for another 30 minutes. Then the cells washed with PBS and stained withZombie Green™ Fixable Viability dye (Biolegend, Cat. No: 423111). Next, the cells were stained with Alexa Fluor® 647 anti-Puromycin Antibody (Biolegend, Cat. No: 381508) using eBioscience™ Foxp3 / Transcription Factor Staining Buffer Set (Invitrogen, Cat. No: 00-5523-00). Incubation with anti-puromycin antibody was done in the permeabilization buffer at 4°C for 1 hour in dark. Flow cytometry was done using BD FACSymphony A5. Geometric mean Fluorescence intensity (gMFI) of Puromycin was measured in all samples using FlowJo-V10. To calculate mitochondrial dependence and glycolytic capacity the following formulas were used:

Mitochondrial dependence (%) = 100* ((DMSO-O)/(DMSO-2DGO))

Glycolytic Capacity (%) = 100-Mitochondrial dependence (%)

### IACS-010759 cytotoxicity measurement

To determine the cytotoxicity of IACS-10759, MOC2 and FaDu cells were cultured in 96 well plates at density of 500 and 2500 cells per well respectively. After 24 hours incubation, IACS was added to the cells at indicated concentrations (Fig 6A) and incubated for 5 days. The medium was replaced with fresh medium contained IACS at day 3. Sulfarhodamine B (SRB) was used on day 5 for cell quantification.

### ATAC-seq

ATAC-seq library prep was done using ATAC-seq Kit (Active Motif, Cat. No: 53150). Briefly, 100,000 SCC1 cells were collected after 72 hours siRNA transfection. Nuclei preparation, tagmentation reaction and purification, PCR amplification of tagmented DNA were done according to manufacturer’s protocol. Libraries size distribution was analyzed using Agilent 2100 bioanalyzer. Libraries were sequenced in 50 bp paired-end reads on Illumina NovaSeq X Plus at MedGenome Inc. Analysis was done using nf-core ATAC-seq pipeline (nf-core/atacseq: 2.1.2) (60). In brief, alignment to human reference genome (GRCh38) was done using Bowtie2 aligner (2.4.4). Narrow peaks were called by MACS2 (2.2.7.1) and differential accessibility analysis was conducted with DESeq2 (1.28.0). Foot printing analysis was performed using TOBIAS 0.13.3.

### ACTL6A CUT&RUN sequencing

DNA prep was done using the ChIC/CUT&RUN Assay Kit (Active Motif, Cat. #53180). Briefly, 500,000 SCC1 cells per reaction were harvested following 72-hour treatment with either DMSO or 1 µM AU-15330. Nuclei isolation, antibody incubation on Concanavalin A–coated magnetic beads, ACTL6A/BAF53A (E3W2A) Rabbit Monoclonal Antibody (#92324) or Rabbit IgG Isotype Control (DA1E) (#66362) binding, pAG-MNase binding, targeted chromatin digestion, and DNA purification were performed according to the manufacturer’s instructions. NGS libraries were prepared using the NEBNext Ultra II DNA Library Prep Kit and sequenced as 50-bp paired-end reads on the Illumina NovaSeq X Plus platform, targeting ∼10 million paired reads (∼20M total) per sample (MedGenome Inc.). Bioinformatics analysis was carried out using nf-core/cutandrun v3.2.2 with default parameters. Reads were aligned to the GRCh38/hg38 human reference genome using Bowtie2 v2.4.4. Peak calling was performed using SEACR v1.3. Signal matrices around peak centers were generated using deepTools v3.3.0 (computeMatrix), followed by heatmap and average profile visualization (plotHeatmap). Differential peak analysis was conducted using DESeq2 v1.28.0, and motif enrichment was performed with HOMER v4.11.

### RNA-seq

Total RNA extraction was done using Trizol/chloroform method. SCC1 cells were collected following transient ACTL6A knockdown in 1.5 ml microcentrifuge tubes. 500 µl Trizol (Invitrogen, Cat. No: 15596026) was added to the cell pellets then phase separation was done by adding 100 µl Chloroform to each tube. The mixture was incubated for 5 minutes at RT followed by centrifugation at 12000g for 15 minutes at 4°c. Top phase was transferred carefully to a new tube and 500 µl Isopropanol was added to precipitate RNA. After 10 minutes incubation at RT, the samples were centrifuged at 12000g for 20 minutes at 4°c. Next, the supernatant was discarded and washing was done with 75 % ice-cold Ethanol twice. RNA was dried at Rt for 10 minutes and suspended with Nuclease free ultrapure water. Libraries were made using Illumina® Stranded mRNA Prep kit (20040534). Sequencing was performed by MedGenome Inc on Illumina NovaSeq X Plus sequencing system. Data analysis was done using nf-core RNA sequencing analysis pipeline (nf-core/rnaseq: version 3.14.0). Differential gene expression analysis was done using DESeq2.

### Immunofluorescence staining of tissue sections (Paraffin embedded)

Fresh cut human HNSCC tissue array slides (HN242c) were purchased from Tissue Array (Derwood, MD). Tissue staining was performed using ACTL6A and p-ERK antibodies. Briefly, slides were baked at 60°C for 2 hours. Then the slides were washed in xylene for 10 minutes (3x) followed by washing in 100%, 95%, 70% ethanol, and DI water for 5 minutes each. Antigen retrieval was done in 10mM Sodium Citrate (PH=6) for 15 minutes in a pressure cooker. Then the slides were cooled down in the same buffer on ice for 30 minutes and washed with 1x PBS twice for 10 minutes. Permeabilization was done in 1x PBS/0.2% gelatin/0.25% Triton buffer for 20 minutes. To block non-specific binding, Tissue sections were incubated in 0.5% BSA (dissolved in the permeabilization buffer) for 1 hour at RT. Primary antibodies (diluted in 1% blocking buffer (for dilutions refer to table 2)) added on tissue sections and incubated in a humidity chamber overnight at 4°C. Next, the slides were washed in PBS twice and permeabilization buffer once for 10 minutes each. Sections were then incubated with secondary antibodies (1 to 400 dilution in 1% permeabilization buffer) for 1 hour at RT. Sections were washed with PBS twice and mounted with ProLong™ Gold Antifade Mountant with DNA Stain DAPI (Invitrogen, Cat No: P36931). Imaging was done using ZEISS LSM700 confocal microscope. Image analysis was performed using Fiji 2.16.0.

### Hypoxia experiments

SCC1 and FaDu cells were cultured in 6-well plate following a transient ACTL6A knockdown and incubated in Xcellbio Avatar bioreactor under either ambient condition or 0.2% hypoxia for 24 hours. Next, the cells were collected for downstream analysis. Proliferation assay under hypoxia was done using Avatar AI system. First, 100 ul DTT (1 Mm) was added to Xcellbio AI 96 well plate (000156) and incubated at 37 ° c for 30 minutes to remove all oxygens and stabilize the electrodes. DTT was removed and wells were washed with 200 µl DPBS. Then the cells were cultured in the plate and incubated under 0.2% hypoxia for 4 days. Data was collected using XcellSoft software v283.

### HRE-luciferase reporter assay

A hypoxia-response-element (HRE) luciferase reporter containing three tandem HRE sites upstream of a minimal TK promoter driving firefly luciferase, with a PGK-driven hygromycin resistance cassette for selection, was used to monitor HIF transcriptional activity (a gift from William Kaelin (RRID: Addgene_118706)). FaDu and SCC1 cells were transduced with reporter lentivirus and selected with 400 µg/ml hygromycin. Reporter-expressing cells were subjected to transient ACTL6A knockdown (siACTL6A) or non-targeting control (siNTC) as described before. 72 hours after transfection, cells were treated with either DMSO or 40 µM T-5224 (Medchemexpress, HY-12270) and incubated under normoxia or 0.2% hypoxia (Xcellbio AVATAR) for 24 hours. Luciferase activity was measured using Pierce™ Firefly Luciferase Glow Assay Kit on a plate reader and reported as relative light units (RLU). n = 3 replicates per condition. To validate AP-1 inhibition, SCC1 and FaDu cells were treated with 50 ng/ml PMA for 30 minutes then with DMSO, T-5224 alone (10, 20, or 40 µM), 50 ng/ml PMA, or PMA combined with T-5224 (10, 20, or 40 µM) for 1 hour, and expression of the AP-1 target gene FOSL1 was measured by qRT-PCR relative to RPL13a.

### Animal studies

All in vivo experiments were done in accordance with the Institutional Animal Care and Use Committee at Stanford University. B6J.Rag2 IL2Rγ double KO (RRID:IMSR_JAX:038282) mice were injected subcutaneously with 500,000 MOC2 shNTC or shActl6a cells. When average tumor size reached 200 mm^3^, IACS-10759 (MedChemExpress, Cat. No.: HY-112037) or vehicle (10% DMSO/40% PEG300/5% Tween-80/45% Saline) was given to mice by oral gavage daily at 10 mg/kg. Tumor width and length were measured by Caliper every 3-4 days and volume was calculated as follows: Tumor volume = Length × width^2^ × 0.5. Mice were euthanized when tumors reached 1000 mm^3^ within one treatment group and tumor tissue was collected for further analysis.

### In vivo HIF reporter imaging and tumor growth

MOC2 cells stably expressing the 3× HRE-luciferase reporter and either control (shNTC) or Actl6a-targeting (shactl6a) shRNA were generated by sequential lentiviral transduction and antibiotic selection. 5 × 10^5 reporter-expressing MOC2 cells were injected subcutaneously into B6J.Rag2 IL2Rγ double KO mice (n = 5-6/group). [MM1.1]Tumor bioluminescence was measured at the indicated time points following intraperitoneal injection of D-luciferin (150 mg/kg) using a SII LagoX imaging instrument (mean radiance (photons/sec/cm2/sr) was quantified over a region of interest using Aura v5.1. Tumor length and width were measured by caliper every 3–4 days and volume was calculated as length × width2 × 0.5. At study endpoint, tumors were harvested and weighed.

### qRT-PCR

RNA isolation was done as described in (*Bulk RNA sequencing*). cDNA was synthesized using HiScript III 1st Strand cDNA Synthesis Kit (Vazyme, Cat. No: R312-01) following manufacturer’s protocol. qPCR reactions were done using Taq Pro Universal SYBR qPCR Master Mix (Vazyme, Cat. No: Q712-02) in QuantStudio 7 Pro Real-Time PCR System. mRNA expression was calculated relative to RPL13a. Primers are shown in table S1.

### Western blot

Cells were washed with cold PBS and lysed using RIPA lysis buffer (Miliporesigma: 20108) supplemented with phosphatase/protease inhibitors cocktail for 30 minutes on ice. Then the lysate was centrifuged at 12000 g for 10 minutes at 4°c. Supernatant was collected and protein concentration was measured using Pierce™ BCA Protein Assay Kit (Thermo Scientific: 23227). 20 µg protein was boiled with Laemmli SDS sample buffer for 5 minutes at 95°c and electrophoresis was done using Novex™ Tris-Glycine Mini Protein Gels, 4–12% (Invitrogen™: XP04205BOX). Transfer was performed on iBlot 3 Western Blot Transfer System. Then the membrane was incubated in Intercept® (PBS) Blocking Buffer (LI-COR: 927-70001) at RT for 1 hour followed by overnight incubation in primary antibody at 4°c. the next day, membrane was washed in TBST buffer (Tris buffered saline + 0.1% Tween 20) three times for 10 minutes followed by incubation in secondary antibody at RT for 1 hour. Membrane imaging and analysis was done using Odyssey CLx Imager and image studio software respectively.

### Co-immunoprecipitation

Co-immunoprecipitation was performed using the Universal Magnetic Co-IP Kit (Active Motif, Cat. No. 54002) according to the manufacturer’s instructions. Briefly, whole-cell extracts were prepared from FaDu and SCC1 cells using the kit’s extraction reagents, with all buffers supplemented with the provided protease, phosphatase, and deacetylase inhibitors and PMSF. Protein concentration was determined by BCA assay. For each reaction, 2 mg of extract was diluted in the kit Co-IP/Wash Buffer and incubated with anti c-Fos antibody (Cell Signaling Technology: 31254s) or an equal amount of normal rabbit IgG as a negative control, together with the kit’s Protein G magnetic beads, rotating end-over-end for 4 hours at 4°C. Bead-bound complexes were collected on the magnetic stand and washed 6 times with Co-IP/Wash Buffer. Immunoprecipitated proteins were eluted by boiling in Laemmli SDS sample buffer, resolved by SDS-PAGE alongside 5% of the input extract, and immunoblotted for c-Fos and ACTL6A as described in the “Western blot “methods

### DepMap studies

We used the Dependency Map Project (DepMap) repository of cell line encyclopedia to search for oral and larynx SCC subsites, which are generally HPV (-). There were 62 cell lines (table S2) for which RNA-seq data was downloaded. Expression data for ACTL6A, glycolysis genes, and Ras genes was exported to GraphPad and a Spearman r co-efficient and p-value was calculated.

### TCGA analyses

Gene expression data for the TCGA cohorts was obtained from the UCSC Xena repository as log2(RSEM+1)-normalized values, restricted to primary tumor samples. For each cohort, a per-sample Buffa hypoxia score was calculated as the mean of the across-sample z-scores of the 36 genes in the Buffa hypoxia signature (ACOT7, ADM, BNIP3, CORO1C, DDIT4, ESRP1, KIF20A, LDHA, LRRC42, MRGBP, MRPL13, NDRG1, P4HA1, PGK1, PNP, PSRC1, SHCBP1, SLC25A32, TUBB6, UTP11L, VEGFA, MRPL15, MAP7D1, TUBA1C, SEC61G, CDKN3, ANLN, CHCHD2, KIF4A, MAD2L2, MIF, MRPS17, PSMA7, SLC16A1, SLC2A1, YKT6). AP-1 transcription factor activity was estimated using VIPER with regulons from DoRothEA. Per-sample AP-1 activity was defined as the mean estimated activity of the AP-1 family members JUN, JUNB, JUND, FOS, FRA1 (FOSL1), and FRA2 (FOSL2). Within each cohort, tumors were z-scored for ACTL6A expression and stratified into ACTL6A-high (top tertile) and ACTL6A-low (bottom tertile) groups, with the middle tertile excluded, and the association between AP-1 activity and Buffa hypoxia score was assessed separately within each group. Correlation coefficient r and p value are reported for each group. To test whether ACTL6A modifies the AP-1–hypoxia relationship, a linear interaction model ((Hypoxia_z ∼ (AP1_z + ACTL6A_z) + (AP1_z × ACTL6A_z)) was fit across all samples. Analyses were performed in R (v4.5) using the dplyr, ggplot2, viper, and dorothea packages.

The HALLMARK_KRAS_SIGNALING_UP and HALLMARK_GLYCOLYSIS gene sets were retrieved from the MSigDB hallmark collection using the msigdbr R package and restricted to genes present in the expression matrix. Per-sample enrichment scores for both gene sets were calculated by single-sample gene set enrichment analysis (ssGSEA) using the GSVA package with score normalization enabled, and enrichment scores and ACTL6A expression were converted to z-scores. Within each cohort, tumors were stratified by ACTL6A expression into ACTL6A-high (top tertile) and ACTL6A-low (bottom tertile) groups, with the middle tertile excluded from analysis. The association between KRAS signaling and glycolysis enrichment was assessed separately within each group using Spearman correlation, and ρ and p values are reported for each group. Analyses were performed in R (v4.5) using the dplyr, ggplot2, msigdbr, and GSVA packages.”

For Figure 7E, A Kaplan-Meier survival curve was generated after stratifying patients by median glycolysis enrichment and a Hazard Ratio and log rank test were generated for statistical analysis in GraphPad.

### CosMx workflow

Single-cell spatial transcriptomics was performed using the CosMx spatial molecular imager and the Human 1K panel containing a standard 1000 probe dataset without modifications according to manufacturer’s protocol (Bruker Spatial Biology). This was performed on a TMA containing adjacent cores from the same samples used for immunofluorescence analysis of ACTL6A and p-ERK expression described above. Raw data were processed on the AtoMx Spatial Informatics Platform and exported as flat files comprising a per-transcript table, FOV position metadata, cell polygon metadata files, and morphology marker images. All downstream analysis was performed in R (v4.5).

Quality control was performed at the slide, FOV, and gene level prior to segmentation using the raw transcript table. Cells were segmented directly from transcript coordinates using Proseg and assembled into the Seurat object. Distributions of detected features and total transcript counts were inspected per sample and cells were retained if they expressed >20 features and >20 total transcripts. Cells above the 99^th^ percentile of total count distribution were removed to reduce segmentation artifacts. Negative control and system control probes were summed per cell and cells in which their percentage exceeded 20% were removed. Filtered count matrices were then normalized using SCTransform to reduce artifact related to transcript abundance per cell.

A reference-based strategy was used for cell type annotation. The scRNA-seq dataset of HNSCC tumors from Choi et al. (GSE 181919) was obtained and processed independently. Cells were clustered de novo and manually annotated based on canonical marker expression from the reference dataset into major lineages. Annotations were projected onto the CosMx dataset using SingleR. Transferred labels were validated by computing differential expression of canonical markers within each lineage and mapping annotations back to tissue coordinates using SpatialDimPlot. FOV and core-level data were manually mapped back onto the TMA layout and integrated into the Seurat file.

### Spatial transcriptomics analysis

Analysis was restricted to 7 oral cavity and larynx SCCs within the TMA. Three tumors were excluded from analysis due to sinonasal subsite (n =3) along with two normal mucosal samples. An ACTL6A-dependent gene score was derived from 265 genes with decreased expression and chromatin accessibility on RNA-seq and ATAC-seq, respectively, after ACTL6A depletion and direct ACTL6A occupancy on CUT&RUN based on an adjusted p-value < 0.01. Of these genes, 29 were represented in the CosMx panel (Table S4). Normalized SCT expression was used to generate a per-malignant cell scaled-mean ACTL6A activity score. Each of the 29 genes was standardized across all 70,736 malignant cells, and a per-cell ACTL6A activity score was computed as the unweighted mean of the 29 standardized values. This score was checked relative to rank-based scoring on RNA counts using UCell for validity. Hallmark Hypoxia and KRAS Up gene sets were obtained and a similar process used to generate per-malignant cell activity scores after excluding genes in the ACTL6A activity dataset.

FOV-level data was extracted and all FOVs containing >30 malignant cells were included. A linear mixed-effects model with ACTL6A activity score as a fixed effect and tumor and core as intercepts was used for statistical comparisons between ACTL6A activity scores and Hypoxia and KRAS signaling scores. FOV-level pseudobulk profiles were used for GSEA of ACTL6A activity. Hallmark gene sets were obtained using msigdbr and the 29 genes within the ACTL6A activity gene set were excluded. fgseaMultilevel was used to associate pathways with ACTL6A activity scores. Benjamini–Hochberg correction was used for statistical comparison.

Spatial neighborhoods were calculated separately within each tissue core. For every malignant cell, endothelial cells and fibroblasts within 50- and 100-µm radii were identified from cell-centroid coordinates using fixed-radius nearest-neighbor searches implemented with dbscan::frNN. The number of endothelial or fibroblast neighbors was recorded for each malignant cell at each radius. Neighborhood measurements were averaged across all malignant cells within each FOV. Mean ACTL6A activity and mean neighborhood measurements were standardized separately within each tumor. A similar linear mixed-effects model as previously described was used to associate ACTL6A activity score with endothelial and fibroblast neighbors.

### Statistical analysis

Statistical analysis for cell phenotypic assays, including metabolism and mitochondrial function, the *in vitro* tumor model, and TCGA Kaplan Meier curve was performed in GraphPad Prism. Data were analyzed using Student’s t-test for pairwise comparison and one- or two-way ANOVA for multiple comparisons. A log rank test was used for survival analysis. Data are presented as mean ± SEM and p-values are noted in the figures and text. GSVA enrichment scores for TCGA data were generated in R (v4.5). RNA-seq, ATAC-seq, and CUT&RUN sequencing data was aligned and analyzed using corresponding nf-core pipelines in Python (v3.14, Python Software Foundation). Sample size (n) is noted in figure legends throughout the text.

