## Supplementary data for "ACTL6A regulates the Warburg effect through coordinated activation of AP-1 signaling in head and neck squamous cell carcinoma"

**Supplementary table 1.** PCR primer and RNA knockdown sequences

| qRT-PCR Primers | Sequence (5' to 3') |
|---|---|
| KRAS F human | TGAGAGAGATCCGACAATACAGAT |
| KRAS R human | GCATCATCAACACCCAGATTAC |
| HRAS F human | ACGCACTGTGGAATCTCGGCAG |
| HRAS R human | TCACGCACCAACGTGTAGAAGG |
| NRAS F human | GAAACCTCAGCCAAGACCAGAC |
| NRAS R human | GGCAATCCCATACAACCCTGAG |
| RPL13A F human | CTCAAGGTGTTTGACGGCATCC |
| RPL13A R human | TACTTCCAGCCAACCTCGTGAG |
| **Mouse shRNAs** | **Gene target sequence** |
| Actl6a | CAAGTTACTGCCTATGCTT |
| | CATACAAGATGCATGTCAA |

**Supplementary table 2.** HPV negative DepMap cell lines

| **Oral and Larynx SCC cell lines** | |
|---|---|
| SCC9 | UPCISCC152 |
| SCC25 | UPCISCC154 |
| BICR31 | BICR78 |
| SCC4 | H103 |
| SCC15 | H157 |
| BICR6 | BICR3 |
| HSC2 | H314 |
| SNU46 | H357 |
| BICR16 | H376 |
| CAL33 | H413 |
| HSC4 | KON |
| BHY | KOSC2 |
| SNU1076 | OSC19 |
| PECAPJ34 | OSC20 |
| SNU1041 | UPCISCC026 |
| PECAPJ15 | UPCISCC029A |
| YD8 | UPCISCC040 |
| SNU1066 | UPCISCC072 |
| SNU899 | UPCISCC074 |
| SNU1214 | UPCISCC099 |
| YD10B | UPCISCC111 |
| PECAPJ41 | UPCISCC114 |
| PECAPJ49 | UPCISCC116 |
| A253 | UPCISCC131 |
| BICR56 | UPCISCC172 |
| HSC3 | UPCISCC200 |
| BICR22 | SAS |
| CAL27 | CA922 |
| YD15 | HSQ89 |
| FADU | HO1U1 |
| UPCISCC090 | UCSFOT1109 |

**Supplementary table 3.** DepMap ACTL6A-glycolysis gene set correlations.

| Gene | p-value | ρ |
|---|---|---|
| GAPDH | <0.001 | 0.44 |
| HK1 | <0.001 | 0.38 |
| ENO1 | <0.0001 | 0.66 |
| PGAM1 | <0.0001 | 0.56 |
| ALDOA | <0.0001 | 0.60 |
| GPI | <0.0001 | 0.62 |
| PFKM | <0.0001 | 0.50 |
| PGK1 | <0.0001 | 0.52 |
| TPI1 | <0.0001 | 0.67 |

**Supplementary table 4.** Gene sets

| Glycolysis | ACTL6A |
|---|---|
| ALDOA | KRT18 |
| ENO1 | LIF |
| GAPDH | EPHA2 |
| GPI | S100A6 |
| HK1 | IL18 |
| HK2 | RARG |
| LDHA | ETV4 |
| PFKP | ALCAM |
| PGAM1 | CALD1 |
| PGK1 | CDH1 |
| PKM | ZFP36 |
| SLC2A1 | DUSP1 |
| TPI1 | ANXA1 |
| | VEGFA |
| | ATF3 |
| | HSPB1 |
| | CCDC80 |
| | DST |
| | MAPK13 |
| | SERPINH1 |
| | TGFB1 |
| | IL1B |
| | MALAT1 |
| | CLDN4 |
| | CSF3 |
| | TPM2 |
| | ICAM1 |
| | CLCF1 |
| | LGALS1 |

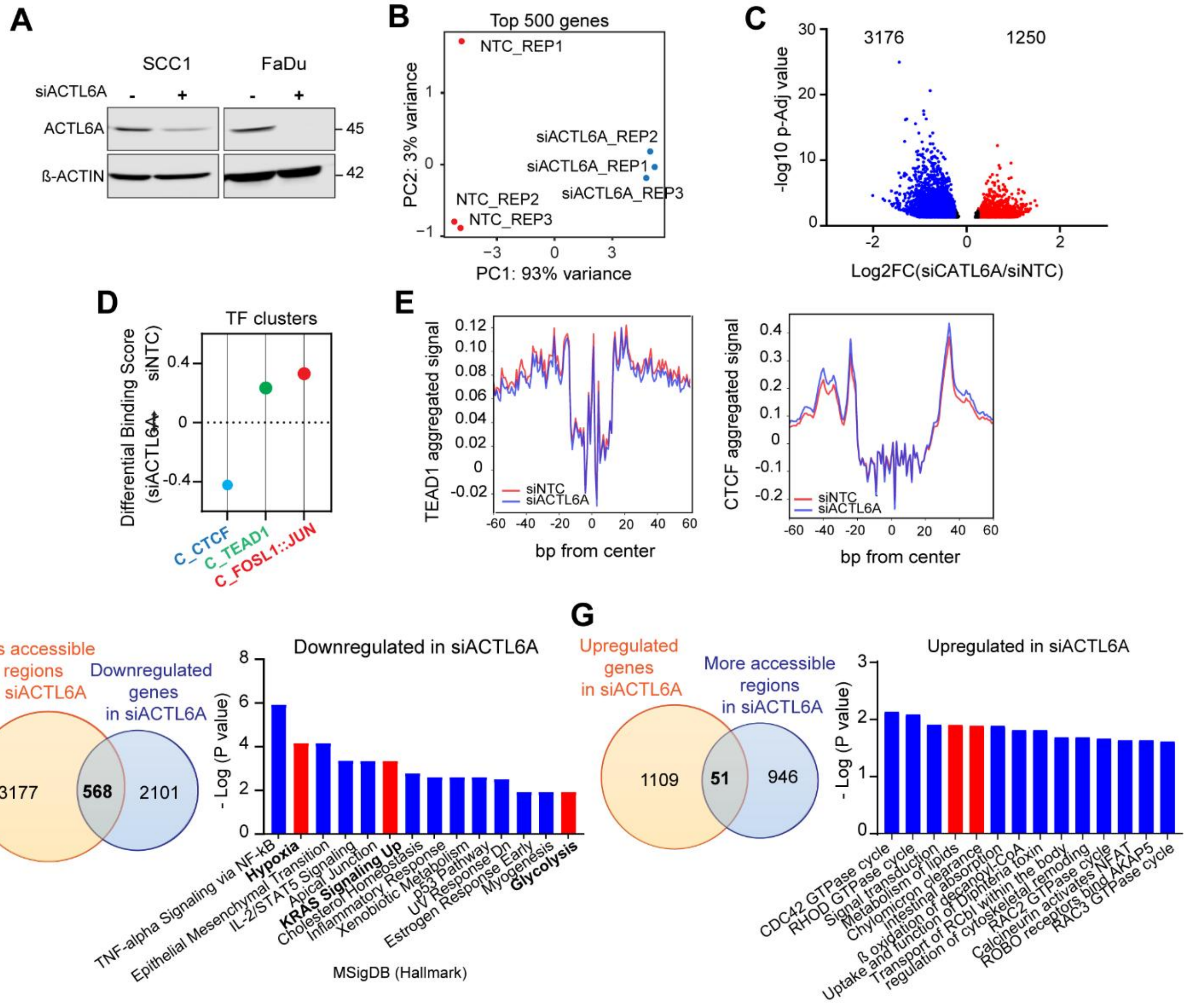


**Supplementary figure 1. ACTL6A depletion alters chromatin accessibility at CTCF, TEAD, and AP-1 transcription factor sites.** (A) Western blot of ACTL6A and β-actin levels in SCC1 and FaDu cells following transient ACTL6A knockdown. (B) Principal component analysis of the top 500 variable genes from SCC1 RNA-seq, showing separation of control (NTC) and siACTL6A replicates (PC1, 93% variance; PC2, 3% variance). (C) Volcano plot of differential chromatin accessibility from SCC1 ATAC-seq , showing 3,176 less accessible (blue) and 1,250 more accessible (red) regions in ACTL6A-depleted cells. (D) Differential binding scores (siACTL6A versus siNTC) for the three transcription factor clusters identified with TOBIAS: C_CTCF, C_TEAD1, and C_FOSL1::JUN. (E) Aggregated footprinting plots centered on TEAD1 (left) and CTCF (right) motifs, comparing SCC1 siNTC and siACTL6A samples. (F) Venn diagram showing overlap between regions with reduced accessibility on ATAC-seq and genes down-regulated on RNA-seq in siACTL6A cells (568 overlapping genes), with MSigDB Hallmark pathway analysis of the overlapping genes (hypoxia, KRAS signaling up, and glycolysis highlighted in red). (G) Venn diagram showing overlap between genes up-regulated on RNA-seq and regions with increased accessibility on ATAC-seq in siACTL6A cells (51 overlapping genes), with corresponding pathway analysis.

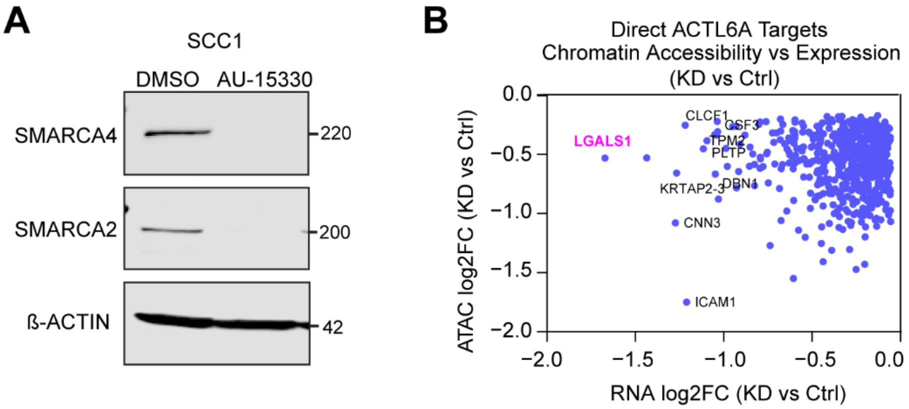


**Supplementary Figure 2. Validation of SWI/SNF degradation and identification of direct ACTL6A targets.** (A) Western blot of SMARCA4, SMARCA2, and β-actin in SCC1 cells following 72-hour treatment with DMSO or 1 μM AU-15330, confirming degradation of both SWI/SNF ATPases. (B) Scatterplot of differential chromatin accessibility (ATAC log2FC) versus differential gene expression (RNA log2FC) for direct ACTL6A targets following ACTL6A knockdown (KD vs. control), identifying LGALS1 as a top hit.

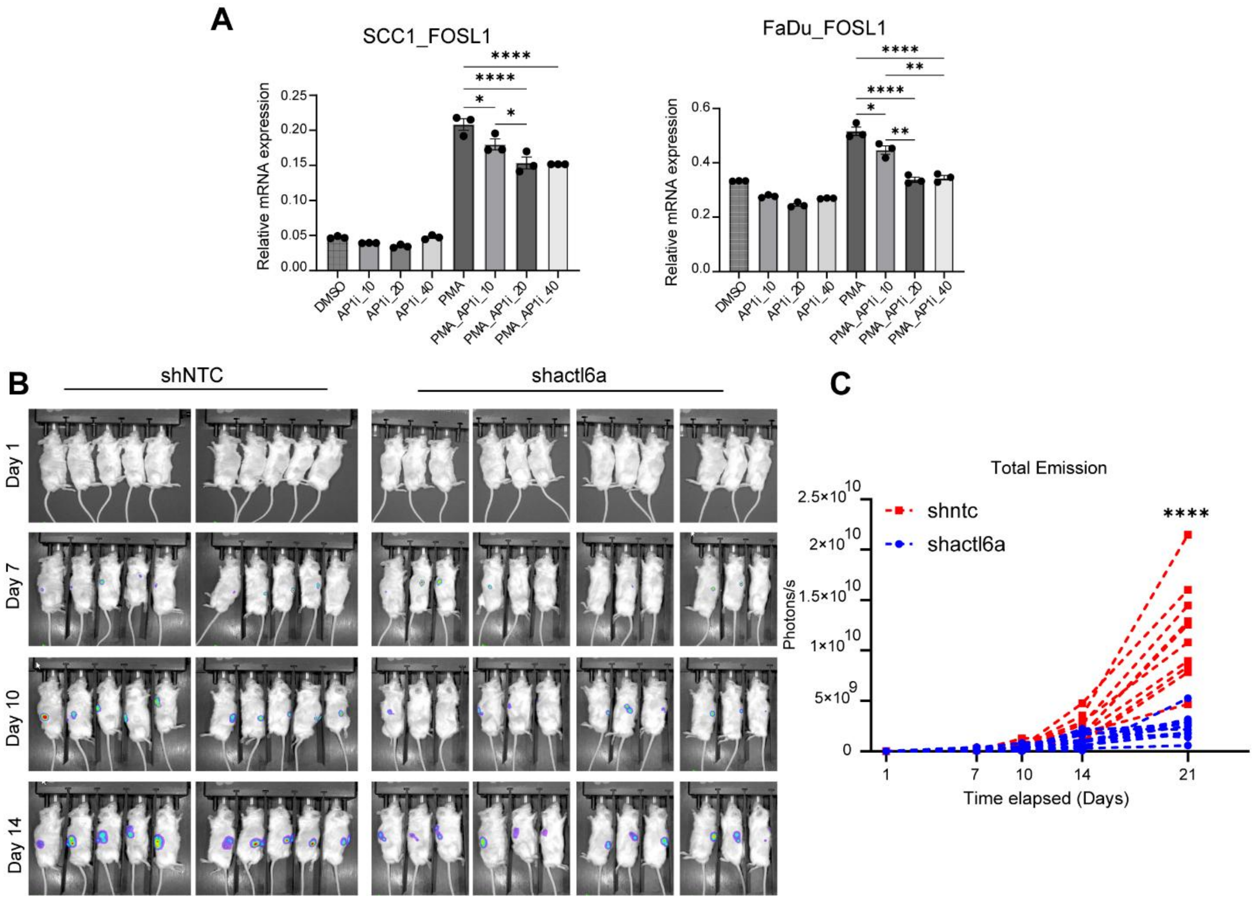


**Supplementary figure 3**. **Validation of AP-1 inhibition by T-5224 and in vivo HIF reporter imaging time course.** (A) qRT-PCR analysis of the AP-1 target gene FOSL1 in SCC1 (left) and FaDu (right) cells treated with DMSO, the AP-1 inhibitor T-5224 alone (10, 20, or 40 µM), the AP-1 activator PMA, or PMA in combination with T-5224 (10, 20, or 40 µM). n = 3. * $p < 0.05$, ** $p < 0.01$, **** $p < 0.0001$, one-way ANOVA followed by Tukey's multiple comparisons test. (B) Bioluminescence imaging time course (days 1, 7, 10, and 14) of mice bearing subcutaneous MOC2 tumors stably expressing the HRE-luciferase reporter with shNTC (left) or shactl6a (right); representative day-21 images are shown in Fig. 3H. (C) Total photon emission (photons/s) of tumors quantified over time for shNTC and shactl6a groups. **** $p < 0.0001$ at day 21, two-way ANOVA followed by Sidak's multiple comparisons test. All data are mean ± SEM.

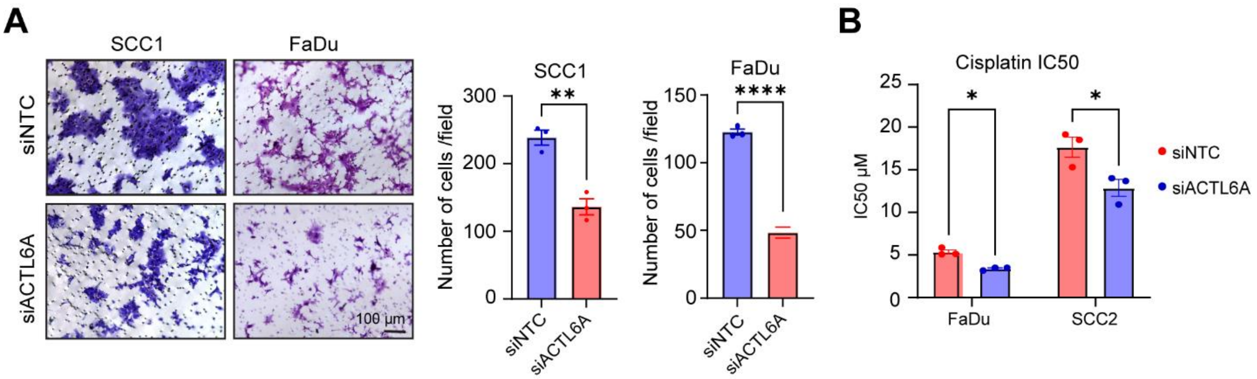


**Supplementary figure 4. ACTL6A depletion reduces cell migration and sensitizes HNSCC cells to cisplatin**. (A) Representative images (left) and quantitation (right) of SCC1 and FaDu cells following transient ACTL6A knockdown after 24-hour incubation in a fibronectin-coated transwell membrane (n = 3). ** $p < 0.01$, **** $p < 0.0001$, unpaired t test. Scale bar: 100 µm. (B) IC50 of cisplatin treatment following transient ACTL6A knockdown in the indicated cells after 72-hour drug incubation (n = 3)., * $p < 0.05$, multiple unpaired t tests. All data are mean ± SEM

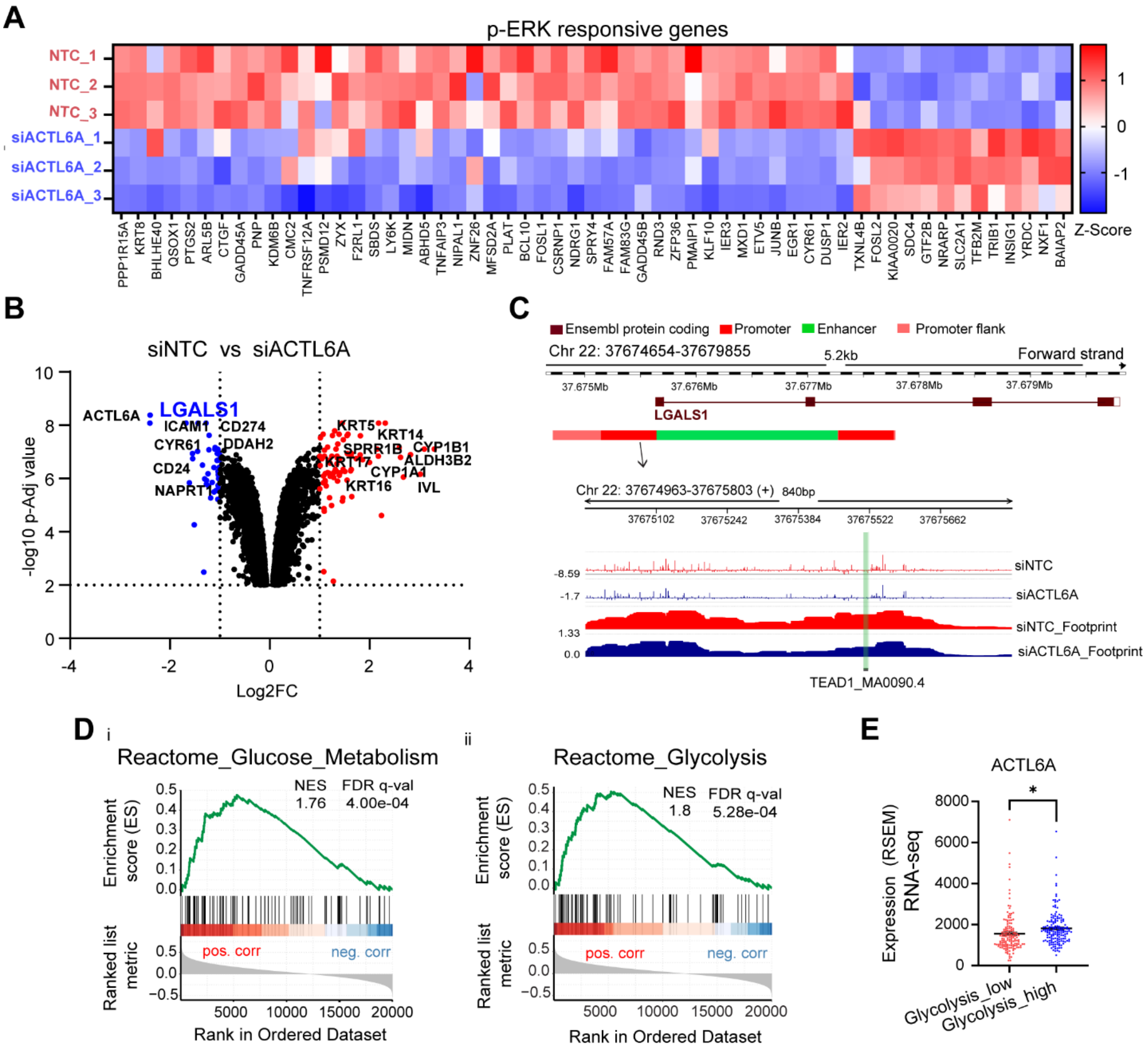


**Supplementary figure 5. ACTL6A regulates p-ERK responsive genes and LGALS1 and is associated with glycolytic gene set enrichment in HNSCC tumors.** (A) Heatmap showing expression of late p-ERK responsive genes using Z-scores generated from SCC1 RNA-seq data for siACTL6A and control replicates. Adj p-value < 0.01. (B) Volcano plot of differentially expressed genes in SCC1 cells following transient ACTL6A knockdown (siNTC versus siACTL6A); significantly down-regulated (blue) and up-regulated (red) genes are labeled, with LGALS1 among the most significantly down-regulated genes. (C) Genomic tracks of the LGALS1 locus (Chr22:37,674,654–37,679,855) showing corrected ATAC-seq peaks and TOBIAS footprints from SCC1 siNTC and siACTL6A cells at a TEAD1 motif (MA0090.4) within the LGALS1 promoter. (D) Enrichment plots for glucose metabolism (i) and glycolysis (ii) Reactome gene sets showing correlation with ACTL6A expression in TCGA HNSCC patients. NES: normalized enrichment score; FDR: false discovery rate. (E) ACTL6A expression (RSEM) in TCGA HNSCC tumors stratified by glycolysis enrichment (low versus high). * $p < 0.05$.

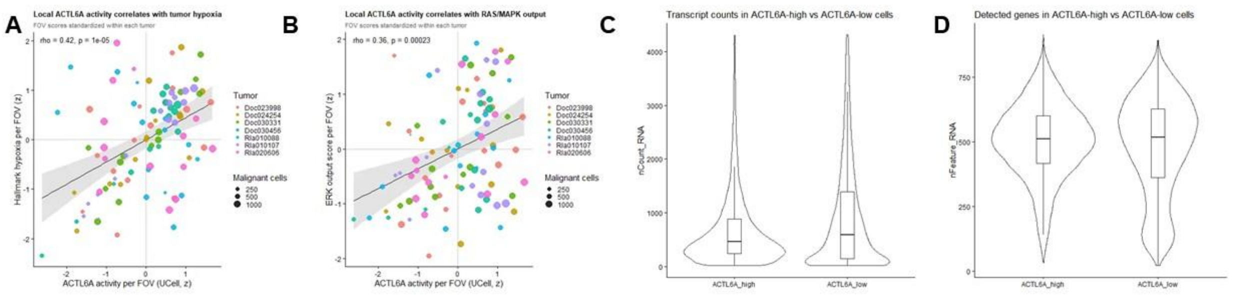


**Supplementary figure 6. Scoring and transcript level validation.** Validation of ACTL6A activity signature using UCell-based scoring showed similar significant FOV-level correlations between ACTL6A activity and (A) Hallmark hypoxia (rho = 0.42, $p < 1x10^{-5}$) and (C) Hallmark KRAS signaling-up (rho = 0.36, $p = 2.3x10^{-4}$) programs. Each point represents one FOV (n = 102 FOVs) with a mean program score across malignant cells within each FOV. Regression lines show linear fits with 95% confidence intervals. (C) Transcript counts and (D) detect genes in malignant cells represented as violin plots with highest quartile (left) and lowest quartile (right) ACTL6A activity scores were not significantly different between groups.

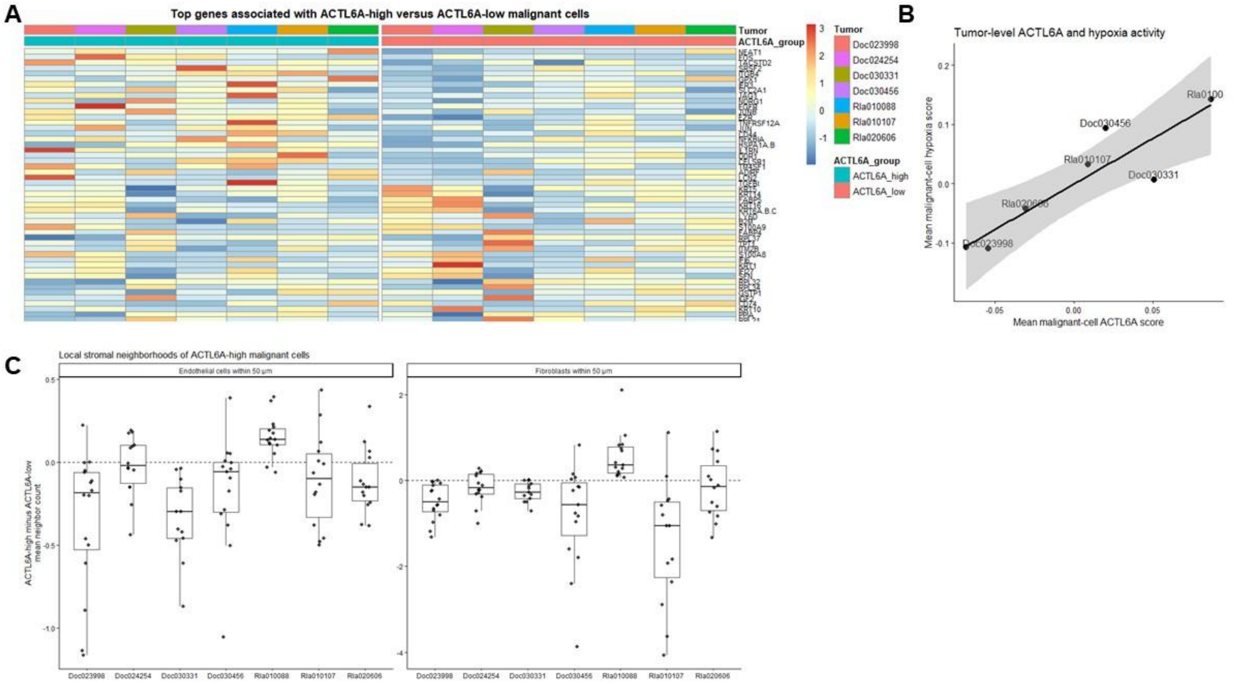


**Supplementary figure 7. Tumor-level ACTL6A activity scores.** (A) Malignant cells were stratified into top quartile (left, blue) and bottom quartile (right, red). A heatmap was generated to visualize the top 25 most up-regulated (top) and top 25 most down-regulated (bottom) genes in ACTL6A activity upper quartile cells relative to the bottom quartile showing tumor level variation. The 29 genes in the ACTL6A activity gene set were excluded. (B) Tumor-level correlation between within-tumor ACTL6A activity score and Hallmark hypoxia scores shows a positive correlation consistent with the FOV-level effect. (C) Tumor-level neighborhood analysis comparing difference in endothelial cell (left) and fibroblast (right) neighbors between upper ACTL6A score quartile and lower ACTL6A quartile. There is a significant decrease in neighbors in 5 of 7 tumors for each cell subset by ACTL6A activity score stratification.

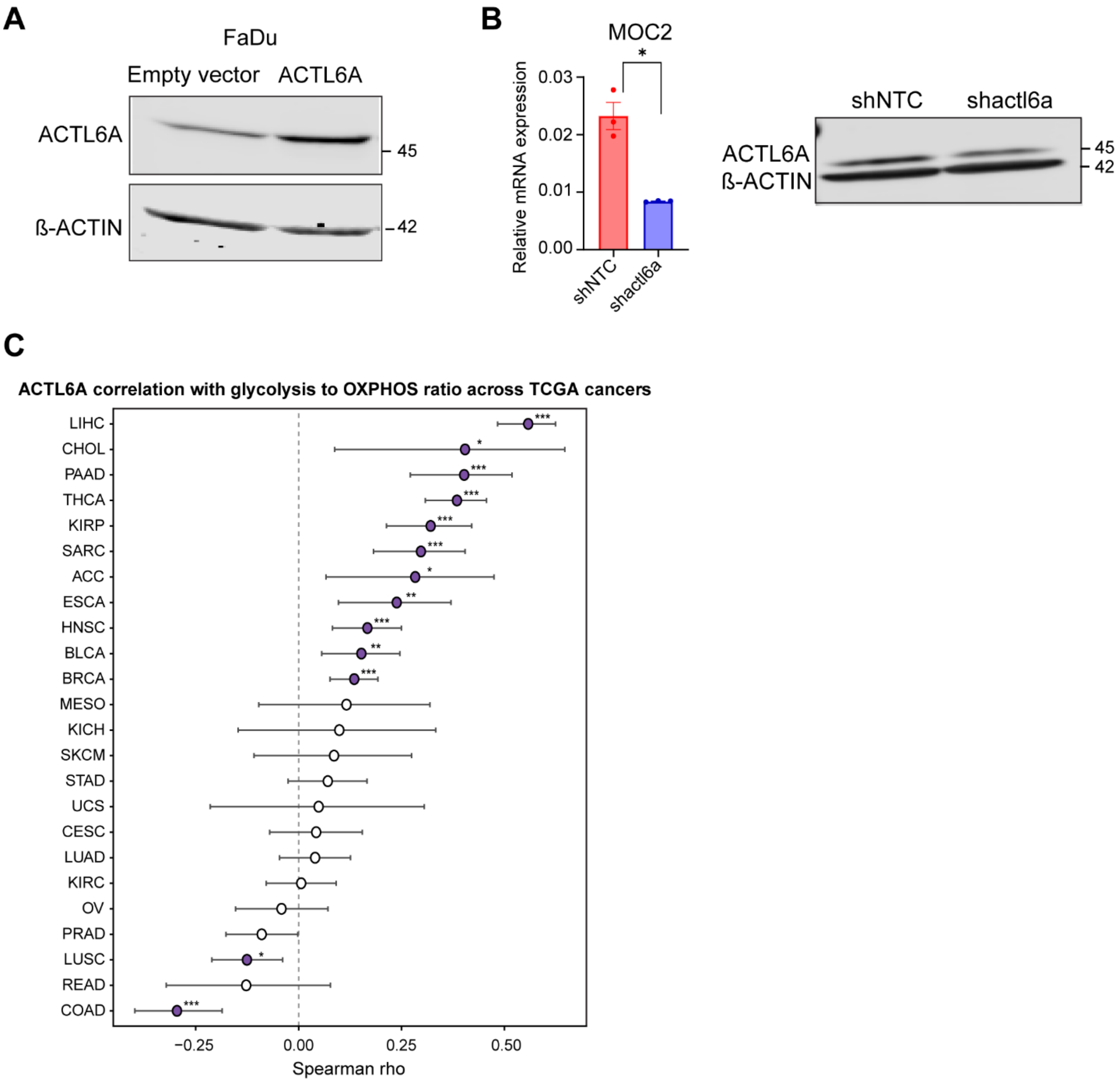


**Supplementary figure 8. *In vivo* MOC2 treatment experiment validation.** (A) Representative Western blot images of β-actin and ACTL6A in FaDu cells after transduction with ACTL6A expressing lentiviral particles. (B) Stable knockdown of MOC2 cells using shRNA against ACTL6A showing RNA levels by RT-qPCR (left) and protein levels by Western blot (right). (C) Forest plot showing Spearman ρ (points) with 95% confidence intervals (horizontal bars) for the correlation between ACTL6A expression and the glycolysis-to-OXPHOS ratio across 24 TCGA cancer types, ordered by effect size. Filled points represents statistically significant correlations and open points represents non-significant correlation. * $p < 0.05$, ** $p < 0.01$, *** $p < 0.001$.
